# Spatiotemporal refinement of signal flow through association cortex during learning

**DOI:** 10.1101/732115

**Authors:** Ariel Gilad, Fritjof Helmchen

## Abstract

Association areas in neocortex encode novel stimulus-outcome relationships but the principles of their engagement during task learning remain elusive. Using chronic wide-field calcium imaging we reveal two phases of spatiotemporal refinement of layer 2/3 cortical activity in mice learning whisker-based texture discrimination. Even before mice reach learning threshold, association cortex—including rostro-lateral (RL), posteromedial (PM), and retrosplenial dorsal (RD) areas—is generally suppressed early during trials (between auditory start cue and whisker-texture touch). As learning proceeds, a spatiotemporal activation sequence builds up, spreading from auditory areas to RL immediately before texture touch (whereas PM and RD remain suppressed) and continuing into barrel cortex, which eventually efficiently discriminates between textures. Additional correlation analysis substantiates this diverging learning-related refinement within association cortex. Our results indicate that a pre-learning phase of general suppression in association cortex precedes a learning-related phase of task-specific signal flow enhancement.

## Introduction

The neocortex dynamically changes when we learn new tasks. Learning to discriminate between different stimuli, e.g. visual stimuli or texture touches, leads to changes in the respective primary sensory areas, i.e., primary visual cortex and barrel cortex (BC)^1–9^. Specifically, neural responses in these areas are enhanced in expert subjects compared to naïve subjects with increased discrimination power to distinguish between stimuli^2,7,10,11^. But what cortical processes during task learning set the ground for this enhancement in primary sensory areas? In other words, are there cortical changes that precede this discrimination, both before the stimulus presentation and also before gaining expertise? Higher-order areas, e.g. retrosplenial cortex and secondary motor cortex, have been shown to mediate learning-induced cortical modulation via top-down effects^4,12,13^. It remains largely unknown, however, how spatiotemporal cortical dynamics reorganizes during learning so that the animal can solve a specific task. Three relevant dimensions to be considered are (1) the large-scale spatial dimension across multiple cortical areas, (2) the temporal dimension relating to the seconds-long duration of individual trials, and (3) the temporal dimension spanning the entire training and learning time course across days.

In terms of the spatial dimension, we have previously applied wide-field calcium imaging to measure large-scale cortical dynamics while mice used their whiskers to discriminate between two textures in a go/no-go task^11^. We found that expert mice displayed enhanced activity for the rewarded go-texture in BC, secondary somatosensory cortex (S2), and rostro-lateral cortex (RL). RL is part of the posterior parietal cortex (PPC)^14,15^, which is part of the higher-order areas in posterior cortex that cluster around the primary visual cortex. These associational areas, and especially PPC, are traditionally thought to represent associations between stimuli and stimulus-outcome sequences: they play a pivotal role in cross-modal sensory integration^16–20^ and hold history-dependent information^21–24^. Nevertheless, the interaction between different association areas and their relationship to learning has not been shown.

The second dimension relates to the temporal sequence of events within an individual trial lasting several seconds. Whereas most studies focus on the time period when the relevant, to-be-learned stimulus is presented to the subject^2–7,13^, little is known about cortical dynamics before the stimulus occurs. For example, when presented with a cue indicating the start of the trial, it is unclear which areas are involved in developing preparatory and anticipatory activity throughout learning^2^. Are there association areas that may show enhanced activity just before a task-relevant stimulus is sensed?

The third dimension relates to the temporal scale across learning, involving many hundreds of trials and lasting several days. Most studies either compare between expert and naïve mice^1–3,5,13,25^ (i.e. two discrete time points) or sample learning on a per-day manner (i.e. 3-8 time points^4,6,7^; but see^9,26^). This relatively low sampling frequency precludes studying the trial-by-trial development of learning, which in some animals can occur within a relatively short period. For example, it is possible that some cortical areas display changes before learning occurs whereas other areas might change when task performance actually improves^9,10,26^.

To study spatiotemporal cortical dynamics during learning we here performed wide-field calcium imaging across a large part of the dorsal cortex while mice learned to use their whiskers to discriminate between two textures that were presented after an initial auditory cue signaling the start of the trial. We measured layer 2/3 activity in 25 cortical regions trial-by-trial continuously during task training across several days. We find learning-related cortical changes—especially in posterior association areas—that can be divided into two phases: First, a pre-learning phase, in which several association areas are suppressed, followed secondly by an enhancement of a specific task-related cortical activation sequence which occurs in parallel with increased task proficiency.

## Results

### Texture discrimination learning

To study learning-related changes in both brain activity and behavior we trained transgenic mice expressing GCaMP6f in layer 2/3 (L2/3) excitatory neurons in a head-fixed, whisker-based go/no-go texture discrimination task^27^ (**Fig. 1a**). Using operant conditioning we trained 5 mice to lick upon whisker-touch with a coarse surface texture (sandpaper with grit size P100) and 2 mice to lick for a smooth P1200 texture. The respective other sandpaper type served as no-go stimulus. An auditory tone served as ‘stimulus cue’ signaling the start of the 2-s long texture approach in each trial. In ‘hit’ trials mice were rewarded for correctly licking for the go texture after an additional auditory ‘response cue’, they were punished with white noise for incorrectly licking for the no-go texture during the response period (‘false alarm’ trials, FA), and they were neither rewarded nor punished when they withheld licking for the go and no-go textures (‘correct-rejections’, CR, and ‘Misses’, respectively). The water spout with lick detector always remained within reach at a fixed position. Licking before the response cue was neither rewarded nor punished. As mice learned to discriminate between the two textures, we measured large-scale neocortical L2/3 activity in the hemisphere contralateral to the texture stimulus using wide-field calcium imaging through an intact skull preparation^11,28^ along with concurrent video monitoring of whisking and body movements (Online Methods). In total, we imaged 7 mice throughout 5-11 days (total trial numbers ranging from 2274 to 5626).

**Fig. 1.**
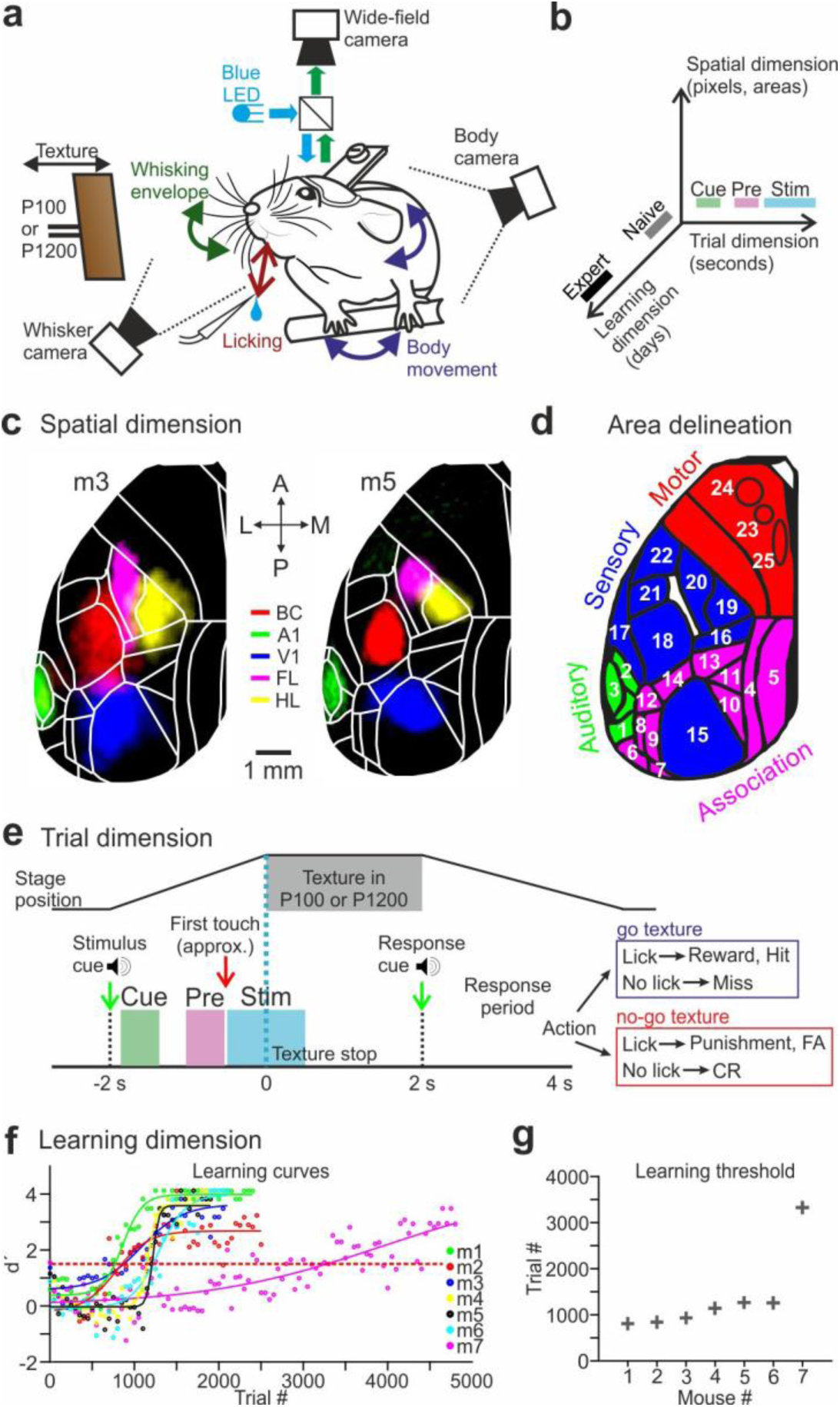
Spatiotemporal dimensions relevant for texture discrimination learning. **a,** Behavioral setup. **b,** Schematic of the three relevant dimensions. **c,** Functional maps for two example mice (m3 and m5) obtained by overlaying sensory-evoked activity maps for different sensory stimuli. Maps are registered to the Allen reference atlas (white outlines). **d,** Area definitions used in this study (see also Supplementary Figure 1). Four rough divisions are auditory areas (green), association areas (pink), sensory areas (visual and somatosensory, blue), and motor areas (red). **e,** Trial structure and possible trial outcomes. After an initial auditory tone (‘stimulus cue’), the P100 or P1200 sandpaper approached and stayed in place for 2 s. At withdrawal start, an additional auditory tone (‘response cue’) signaled that the beginning of the report period. Cue-, pre- and stim-period as analysis windows are marked by different colors. **f,** Performance (d’) for all mice across the entire learning period, fitted with a sigmoid function. Red dashed bar indicates threshold for learning (d’=1.5). **g,** Learning threshold for all mice in ascending order.

We focused our analysis on the three spatiotemporal dimensions mentioned in the introduction (**Fig. 1b**). For analysis of the spatial dimension—covering the entire dorsal cortex— we functionally mapped sensory areas for each mouse during anesthesia. Based on these maps (together with skull coordinates) we registered all wide-field images to the 2D top view of the Allen reference atlas^29^ and defined 25 areas of interest, consolidated in 4 major groups (**Fig. 1c,d** and **Supplementary Fig. 1** with a list of region abbreviations; Online Methods). As one relevant temporal dimension we analyzed signals on the fast time scale of the individual trials (**Fig. 1e**). Here, we were especially interested in the time period before the touch occurred—in addition to the touch-sensation period—because learning-related changes can be expected early in trial time. We defined three time windows during this time period: the ‘*cue-period*’ (0.1 to 0.6 s after the stimulus cue) to capture the responses to the initial tone signaling the trial start; the ‘*pre-period*’ when the texture approaches the whiskers (−1 to −0.5 s relative to the texture stop; mainly before the first whisker-texture touch occurred); and the ‘*stim-period*’ during texture touch (−0.5 to 0.5 s relative to texture stop). We did not include the response period in our analysis because mice licked and moved rigorously during this time period, causing large-scale activity across the cortex that is difficult to interpret (there was no delay period and mice were free to lick outside the response window). The second relevant temporal dimension is the much slower time scale across the days of learning. All mice increased their performance with training (5-11 days; ∼500 trials/day), eventually reaching high discrimination levels as quantified by calculating the d-prime (d’) in a 50- trial bins (**Fig. 1f**; refs. ^11,27^; Online Methods). Performance increased mainly because of an increase in the CR rate (**Supplementary Fig. 2**). In the learning curves we defined the ‘learning threshold’ for reaching expert level as the crossing point at d’=1.5 (in units of ‘trial number’). To compare ‘naïve’ versus ‘expert’ phase we averaged across the first 500 and the last 500 trials, respectively. The fastest learning mouse reached the learning threshold slightly before a thousand trials whereas mouse #7 took a substantially longer time (**Fig. 1g**). Jointly, these definitions of cortical areas, trial periods of interest, and naïve-to-expert learning phases enabled us to reveal key learning-related changes in both behavior and L2/3 activity across the cortex.

### Motor behavior changes during learning

We first quantified changes in motor actions during learning. Mice may start moving more or differently when they begin to associate the go-texture with the upcoming reward. Because movements are accompanied by wide-spread cortical activity^11,30,31^ changes in motor behavior potentially confound the interpretation of learning-related changes of cortical activity. Indeed, once mice reached expert level they indicated their future action before the response cue, by moving their body and by whisking and licking rigorously. To quantify body movements, we detected forelimb and back movements in the body camera video and calculated the movement probability across trial time and across learning (50-trial bins; ref. ^11^, Online Methods). When mice approach and reach their learning threshold, they begin to move their body earlier, clearly before the response cue and reward consumption, resulting in a significantly higher movement probability in the stim-period for the expert compared to the naïve phase (**Fig. 2a,b** and **Supplementary Fig. 3**; p<0.05, n = 7 mice, Wilcoxon signed-rank test). In the pre-period, movement probability was relatively low and did not significantly change from naïve and expert (p=0.47), whereas in the cue-period movement probability showed a significant reduction in expert mice from a relatively low naïve level (p<0.05; Wilcoxon signed-rank test).

**Fig. 2.**
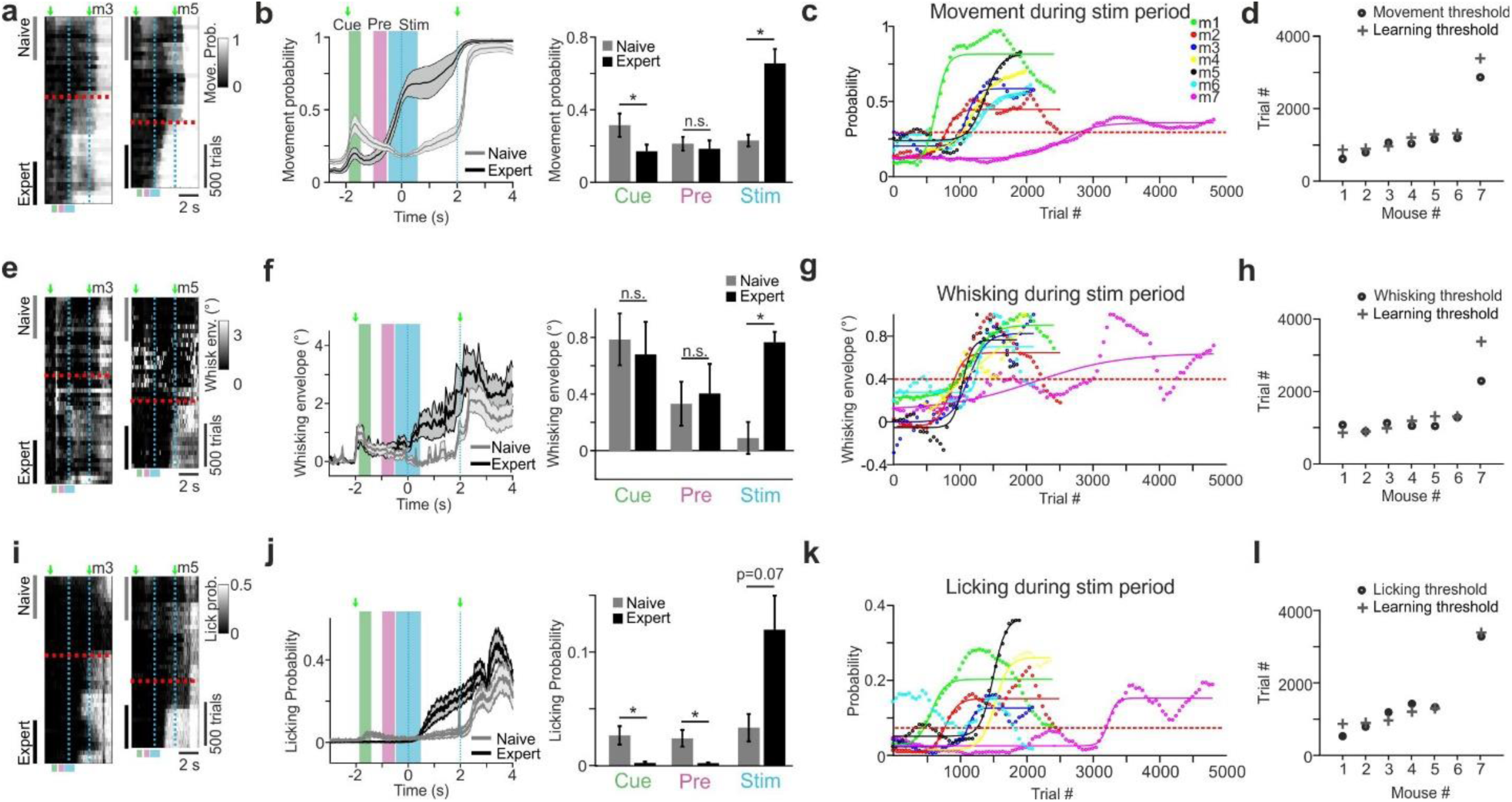
Motor parameters during stim period are associated with learning. **a**, Movement probability for go-trials of two example mice plotted as heat maps along the two temporal dimensions (trial dimension on x-axis; learning dimension on y-axis; 50-trial bins along learning dimension). Red dashed line indicates learning threshold. Cyan dashed lines demarcate ‘texture in’ period. Naïve and expert phases were analyzed in the 500 first and last trials, respectively. **b**, *Left:* Time course of movement probability averaged across all 7 mice during go-trials for naïve phase (grey dashed line) and expert phase (black solid line), respectively. Error shading indicates s.e.m. Green arrows indicate stimulus and response cue. *Right:* Mean movement probability (± s.e.m) in cue-, pre-, and stim-period for naïve and expert phase. n = 7 mice; *p<0.05, n.s. not significant, Wilcoxon signed-rank test. **c**, Movement probability during stim-period across learning for all mice. Each curve was fitted with a sigmoid function. Dashed red line indicates movement threshold (=0.3). **d**, Movement threshold for all mice (circles). For comparison the learning thresholds from Fig. 1g are shown (crosses). **e-h**, Same plots as a-d for whisking behavior. The envelope amplitude of whisking was analyzed. Whisking threshold was defined as 0.4°. **i-l**, Same plots as a-d for licking behavior. The probability of licking was analyzed. Licking threshold was defined as 0.075. Licking threshold could not be detected for mouse #6.

To further understand the relationship between learning and body movement, we calculated the average movement probability during the stim-period for each mouse throughout learning (**Fig. 2c**). By setting a movement threshold of 0.3 (i.e. moving in 30% of the trials) we could accurately predict the learning threshold (**Fig. 2d**; r=0.99, p<0.001 between learning and movement threshold across mice). Thus, although not yet receiving any reward during the stim-period, mice started to move more extensively shortly after texture sensation almost exactly at that time point during training when they learned that the go-texture was associated with a reward.

We performed the same analysis for whisking and licking behavior. On average, the amplitude of the whisking envelope significantly increased with learning in the stim-period (p<0.05, n = 7 mice, Wilcoxon signed-rank test) but showed little change in cue- and pre-period (**Fig. 2e,f**). As for body movements, increased whisking during and following texture touch occurred in parallel to the learning curve and the learning threshold could be well predicted by defining a whisking threshold (**Fig. 2g,h**). With regard to licking behavior during go-trials we found that licking was reduced in expert animals in the cue- and pre-period—highlighting the key requirement of lick suppression for learning—whereas it was enhanced in the stim-period (**Fig. 2k,l**). This increase did not reach significance, presumably because pronounced licking started only at the end of our defined stim-period but was clearly enhanced thereafter in the expert phase. We conclude that mice exhibit consistent learning-related changes in motor behaviors, engaging their body to solve the discrimination task and receive the reward. Once mice learned to discriminate between textures, they initiate various movements during the stim-period whereas they remain relatively quiet before texture touch (i.e. during cue- and pre-period). We therefore first focused on the time period before touch, allowing us to study learning-related changes of cortical processes that lead up to the task-relevant stimulus without the confound of movement-related cortical activity.

### Learning-related changes in cortical activity early during trials

We next analyzed the spatiotemporal dynamics of L2/3 cortical activity across learning, as revealed by wide-field calcium imaging. In the following we present the results for go-trials, i.e. for one type of texture stimulus (hit and miss trials pooled together). We calculated activity maps by averaging ΔF/F signals during the cue-, pre-, and stim-period, respectively, and compared maps obtained during naïve (first 500 trials) and expert (last 500 trials) phase. Maps from two example mice show activation during the cue-period in A1 (also anterior-medial and hindlimb areas) and lower activation in postero-medial (PM) and retrosplenial-dorsal (RD) association areas (**Fig. 3a**). During the pre-period, RL displayed high activation especially in expert mice. During the stim-period, BC displayed the highest activation level. These example maps suggest that a specific sequence of activation emerges during learning, starting from auditory cortex in response to the stimulus cue, followed by RL activation as the texture approaches the whiskers, and continuing to BC activation during touch sensation. This notion of sequential activation is further highlighted by plotting the time course in the expert phase for the mean ΔF/F traces in the regions A1, RL, and BC (**Fig. 3b** for the two example mice; **Supplementary Fig. 5** for all mice).

**Fig. 3.**
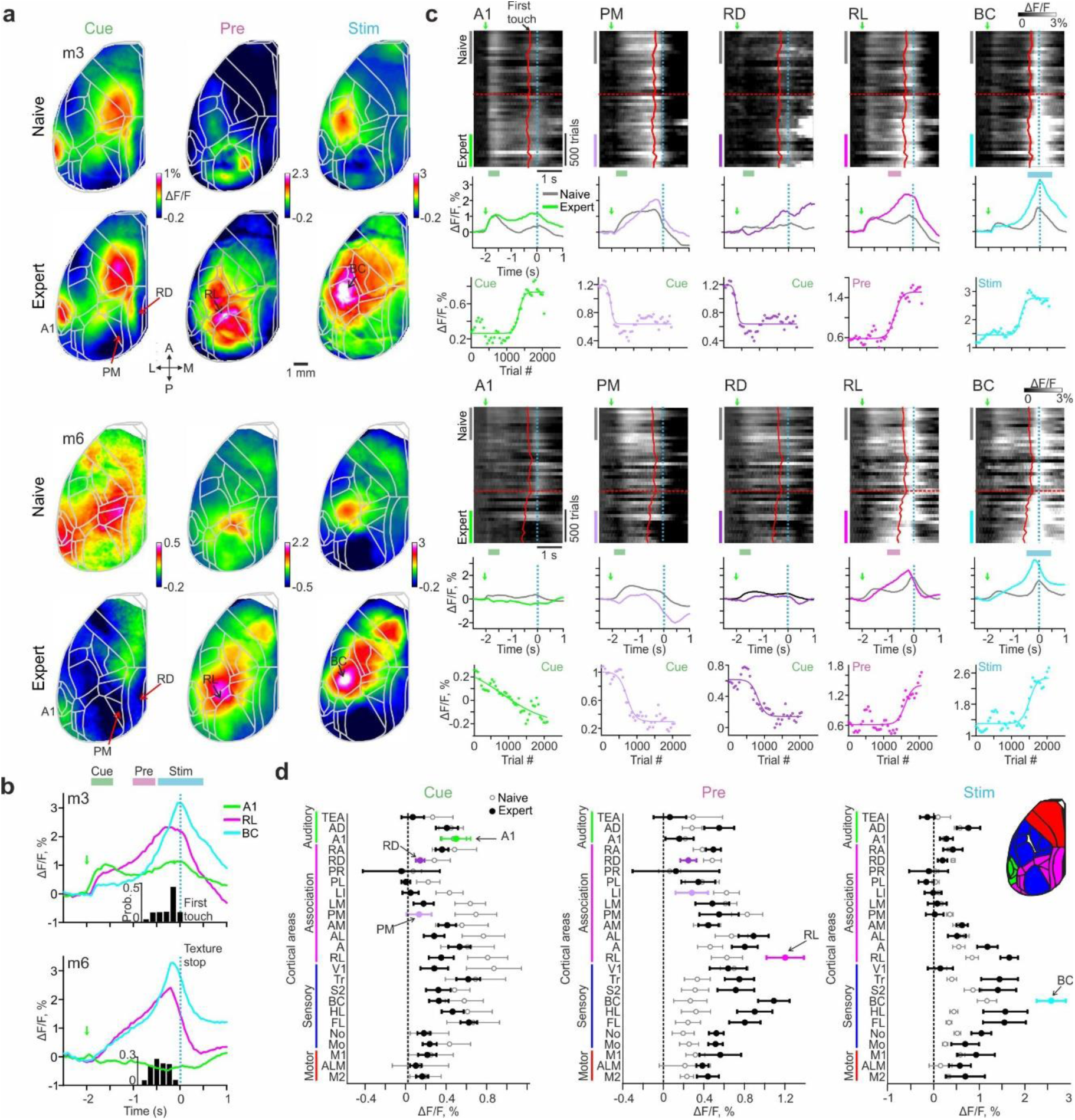
Changes in wide-field calcium signals across cortex during learning. **a**, Example activation maps from two mice averaged during cue-, pre- and stim-period in naïve (top) and expert (bottom) phase. Color scale bar indicates min/max of percent ΔF/F. Overlay of areas in gray for all maps. 5 areas of interest are marked **b,** Expert trial-related responses in A1, RL and BC for the two example mice. Green arrow marks stimulus cue. **c,** Continuation of examples in **a**. *Top:* Population responses plotted for the two temporal scales (trial scale along x-axis; learning scale along y-axis) for five areas of interest (from left to right): A1, PM, RD, RL and BC. Data is binned every 50 trials. Red lines indicate mean first touch of the whiskers on the incoming texture. Dashed red line indicates learning threshold. *Middle:* Trial-related area responses for expert (colored) and naïve (gray). *Bottom:* Mean responses across learning for each area averaged during the specific trial period indicated. Each curve is fitted with a sigmoid function. **d,** Mean activation of all 25 cortical areas in expert (black) and naïve (gray) mice during cue-, pre- and stim-period and grouped into auditory (green), association (pink), sensory (blue) and motor (red) areas (see also inset). Error bars are s.e.m. across mice (n=7).

Next we analyzed in more detail how the cortical activation sequence from cue- to pre- to stim-period changes across learning. Surprisingly, we found learning-related changes during these early trial periods before texture touch. Based on the example activation maps we focused on 5 areas of interest during specific time periods: A1, PM and RD during the cue-period; RL during the pre-period; and BC during the stim-period. For each area we plotted the heat map of trial-related ΔF/F signals across learning (i.e. trial dimension against learning dimension), the comparison of naïve and expert average ΔF/F traces, and the mean ΔF/F responses in the respective trial period across learning (**Fig. 3c**). In the cue-period, A1 activity displayed variable changes during learning, increasing in one mouse while slowly decreasing in the other. We will return to this variability further below. Interestingly, RD and PM showed a suppression of responses across learning for the cue-period. In contrast, in the pre-period RL responses showed consistent enhancement during learning. Finally, BC activity in the stim-period was enhanced in the expert phase. Thus, cortical areas display diverse learning-related changes during specific trial periods, ranging from enhancement (e.g., RL and BC) to suppression (e.g., PM and RD).

We expanded our analysis to all 25 cortical areas by calculating the mean ΔF/F activation for each area during the cue-, pre- and stim-period, averaged across all mice for expert and naïve phase (**Fig. 3d**). During the cue-period, several association areas, including PM and RD, showed reduced activation in expert mice. During the pre-period, expert mice displayed a saliently enhanced activation in RL (actually the largest ΔF/F change for this period) whereas PM and RD maintained lower activation levels. Notably, RL still displayed strong activation during the pre-period when we positioned the texture out of reach of the whiskers in two expert mice so that the relevant stimulus was omitted (**Supplementary Fig. 4**). This finding suggests that RL responses do not directly relate to texture touch *per se* and possibly rather represent the expectation of the upcoming touch. Finally, BC displayed the highest activation in expert mice during the stim-period, along with activation of other sensory and motor areas that presumably relates to the initiation of movements during this period.

### Two phases of cortical changes: wide-spread suppression followed by specific enhancement

Because we imaged large-scale cortical activity throughout the entire learning process, we could relate the time course of learning-related activity changes in the various areas more precisely to the behavioral learning curve. As a first step, we applied sigmoidal fits to the mean ΔF/F changes in the areas of interest selected in Figure 3 and compared the time courses of the normalized curve fits (**Fig. 4a**; for non-normalized traces for all mice see **Supplementary Figure 5**). Surprisingly, activity in PM and RD showed suppression long before the enhancement in BC and RL and even clearly before the behavioral learning threshold was reached. The inflection point where the normalized curve crossed 0.5 occurred significantly earlier for PM and RD than for RL and BC and also significantly preceded the learning threshold (p < 0.05; Wilcoxon signed-rank test; **Fig. 4b** for individual mice and **Fig. 4c** averaged across all mice). Suppression in PM and RD occurred about 500 trials before the mouse reached learning threshold and before the enhancement in RL and BC. In addition, for each of these four areas the inflection point positively correlated with the behavioral learning threshold across mice (r = 0.97. 0.97, 0.79 and 0.79 for BC, RL, PM and RD respectively; p < 0.05). Consequently, the early suppression in PM and RD could predict well when the mouse will learn the task hundreds of trials in advance. In contrast, the inflection points for BC and RL were not significantly different from the learning threshold (p > 0.05; Wilcoxon signed-rank test), implying that they occur rather in parallel with increases in d’ and thus cannot predict when threshold is reached.

**Fig. 4.**
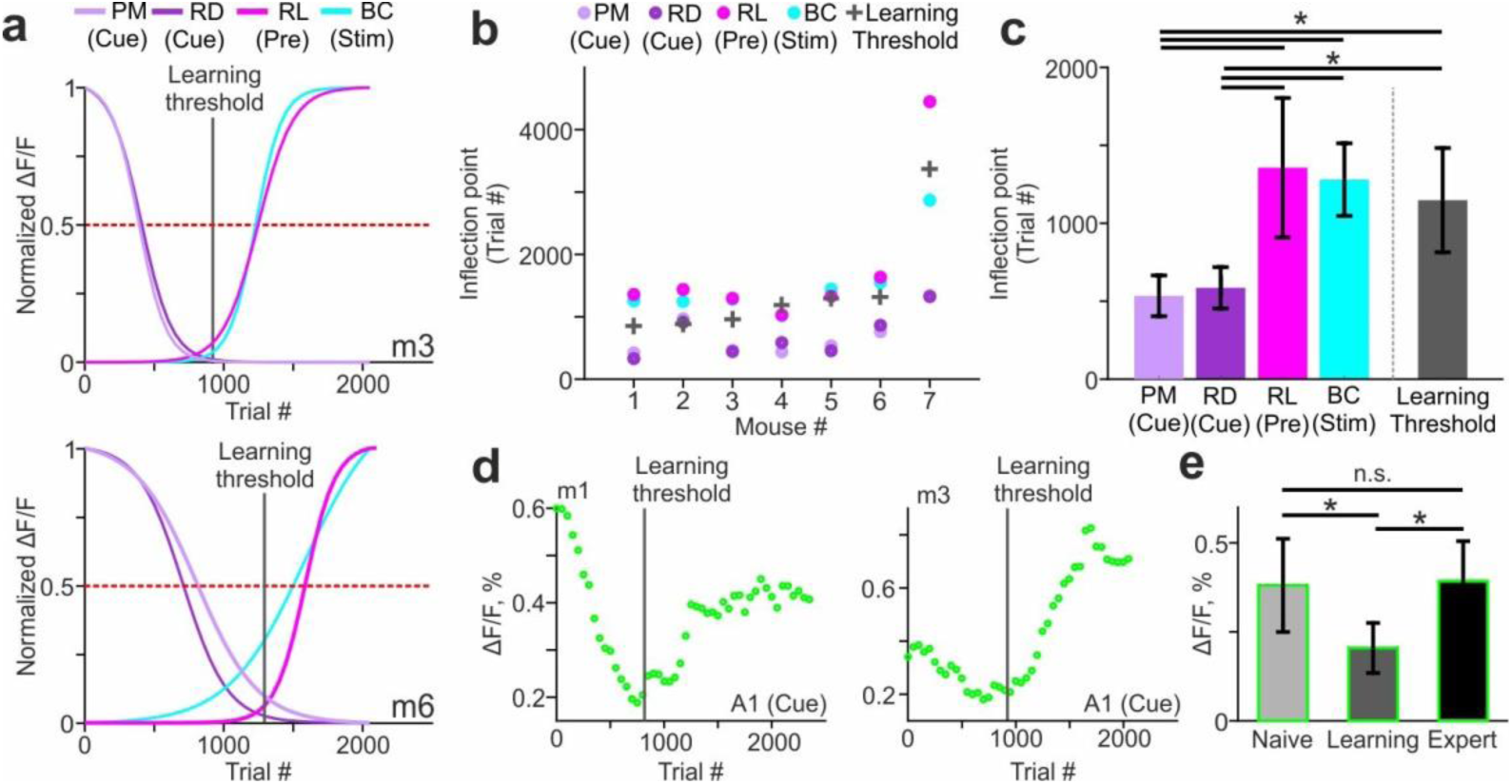
PM and RD suppression occurs before learning and precedes RL and BC enhancement. **a**, Normalized sigmoidal fits to the learning-related mean ΔF/F changes in PM and RD (for cue-period), in RL (for pre-period), and in BC (for stim-period) for two example mice. Horizontal dashed red line indicates 0.5-level for determining steepest points of change. Vertical solid gray line indicates the learning threshold of the mouse. **b,** Steepest points of change for all mice in each of the four areas. Learning threshold is marked with a gray plus sign (similar to Fig. 1g). **c,** Steepest points of change and learning threshold averaged across mice. Error bars are s.e.m. across mice. **d,** Learning-related ΔF/F changes in A1 during the cue-period for two example mice. Vertical gray line indicates learning threshold. **e,** Mean ΔF/F changes in A1 during the cue-period for naïve, learning and expert mice. Error bars are s.e.m. across mice. *p < 0.05; n.s. – not significant; Wilcoxon signed-rank test.

As indicated in Figure 3c the learning-related changes of A1 activity during the cue-period varied widely, with one example mouse displaying mostly enhancement and the other mostly suppression. A closer look reveals that suppression and enhancement were discernible as two consecutive phases in the A1 signals (**Fig. 4d** and **Supplementary Figure 6**). Plotting the cue-period A1 signal changes across learning, together with the respective learning thresholds, we noticed that suppression consistently occurred before mice reached learning threshold whereas enhancement occurred thereafter. The relative amplitude of modulations (suppression or enhancement) varied between mice but A1 calcium signals in the cue-period were significantly lower in amplitude around the time of learning compared to naïve and expert phases (**Fig. 4e**; p < 0.05; Wilcoxon signed-rank test; averaged across ±100 trials around threshold). Thus learning-related changes of activity in cortical areas do not need to be uni-directional, i.e., exclusively decreasing or increasing, they may display mixed effects. Apparently, stimulus-cue induced A1 activity is suppressed early during training before learning, similar to PM and RD, and then dynamically shifts to enhancement after the learning threshold has been reached, similar to RL and BC but to a variable degree.

We wondered whether such two phases of pre-learning suppression and learning-related enhancement are also apparent in other cortical areas and for different trial periods. We therefore applied a two-phase model to all areas by fitting the learning-related ΔF/F signals in cue-, pre- and stim-period with a double sigmoid. This analysis corroborated the concept of two phases of cortical activity changes and substantiated the pre-learning suppression in association areas and later specific enhancement in task-related areas in congruence with learning (**Supplementary Figure 7**).

As an alternative approach to quantify the relationship between the learning curve of the mouse and the learning-related changes in cortical activity, we defined a learning map for the cue-, pre-, and stim-periods by correlating the learning curve (d’ values) with the corresponding time course of the ΔF/F signals for each pixel (averaged over the respective trial period; **Fig. 5a**). The maps for two example mice reveal that several association areas display negative correlation values during the cue-period (**Fig. 5b**), reflecting the predominance of suppression in these areas during the early cue-period. During the pre-period, RL displayed the highest positive correlation whereas PM and RD specifically maintained negative values. Finally, BC displayed the highest correlation during the stim-period. Other sensory and motor areas showed strong correlation, too, presumably reflecting behavior-related neural activity. The divergence of activity patterns across areas during the pre-period (positive correlation with learning in RL and BC vs. low or negative correlations in PM and RD) also became obvious when plotting correlations with the learning curve for each time frame during the trial period (**Fig. 5c**). Pooled across mice correlations between activity and learning were mostly negative in our 5 prime areas during the cue-period, then became significantly positive in RL and BC for the pre-period (while staying significantly negative for PM and RD) and significantly positive in BC for the stim-period (along with RL; **Fig. 5d**; p<0.05; Wilcoxon signed-rank test). Across all 25 areas, many association areas were negatively correlated with d’ values during the cue-period, followed by a spatial refinement during the pre-period with RL displaying positive correlation with learning whereas PM and RD exclusively pertained negative correlations (**Fig. 5e**). As indicated by the learning maps, BC displayed positive correlation for the stim-period as well as most of the sensory and motor areas.

**Fig. 5.**
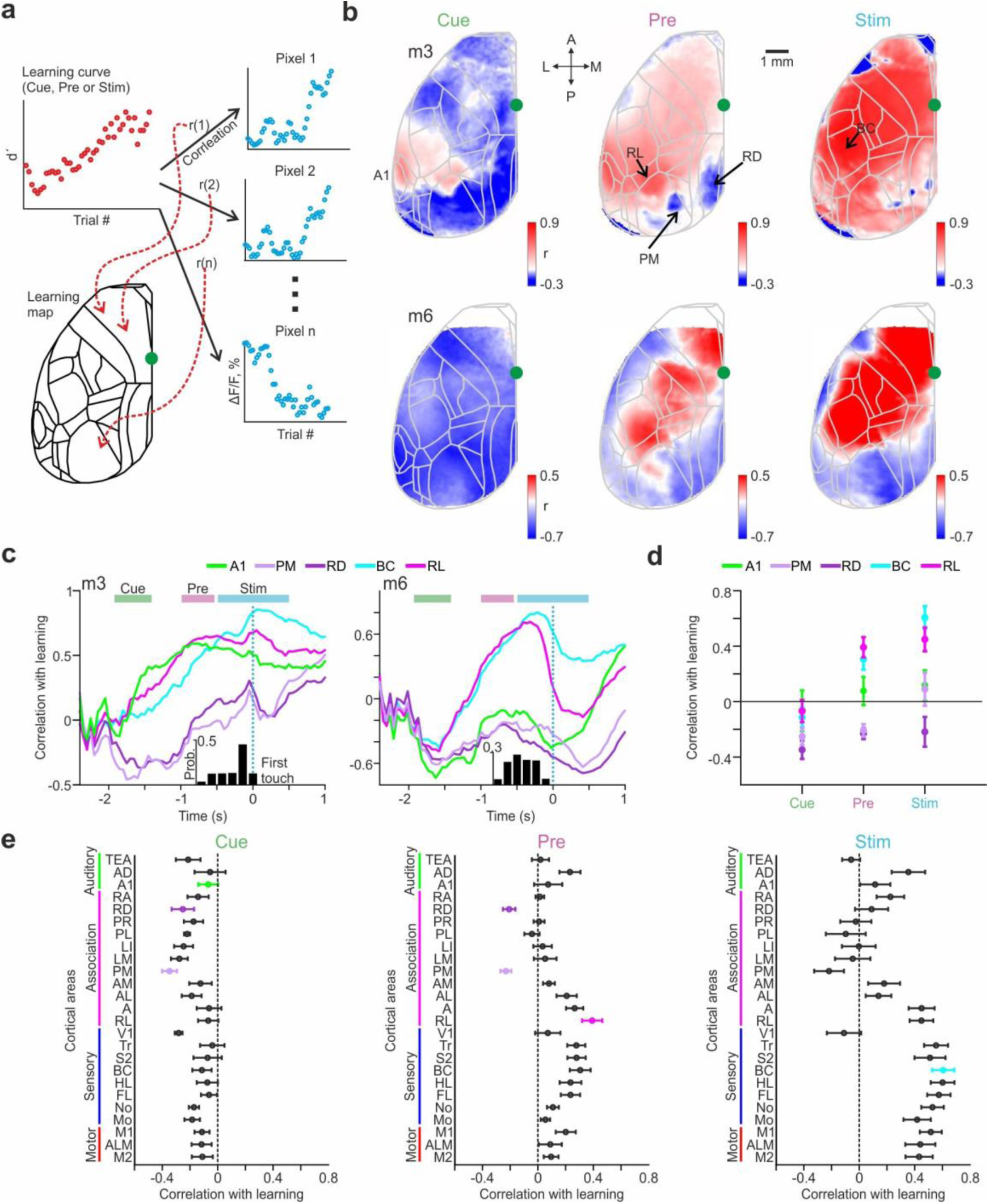
Learning maps reveal dissociation of association areas in relationship to learning. **a**, Schematic illustration for calculating a learning map. Each pixel in the maps reflects the correlation coefficient (r) between the mouse’s learning curve and the curve of learning-related ΔF/F change of the respective pixel. The latter can be averaged across the key trial periods (i.e. cue, pre, or stim) or calculated for each time frame. **b,** Learning maps during cue-, pre-, and stim-periods in two example mice. Color denotes r-values. **c,** Correlation with learning as a function of time for the 5 key areas in two example mice. **d,** Correlation with learning during cue-, pre-, and stim-periods for the 5 areas. Error bars are s.e.m. across mice. **e,** Correlation with learning during cue-, pre-, and stim-periods for all areas. Error bars are s.e.m. across mice.

These findings highlight the spatial refinement that the association areas undergo during learning, especially during the trial period bridging the initial stimulus cue and the arrival of the texture as task-relevant stimulus.

### Seed pixel analysis reveals dissociation within the association network

We next aimed to further quantify the differences and potential interactions among different areas during learning. Similar to the learning maps we calculated ‘seed maps’, for which—instead of correlating ΔF/F signals pixel-wise to the learning curve—we correlated the learning-related ΔF/F changes for all pixels with the reference time course in a ‘seed’ area (**Fig. 6a**). Guided by the learning maps, we first calculated pre-period maps with the association areas RL, PM, or RD as seed areas. The RL seed map revealed a positive correlation of activity in this area with sensory and motor cortices as well as with adjacent association areas (**Fig. 6b**). In contrast, PM and RD seed maps showed high correlations among each other and with their adjacent areas but lower correlations with RL, BC, and most of the other cortical areas (**Fig. 6c**; pooled across all mice).

**Fig. 6.**
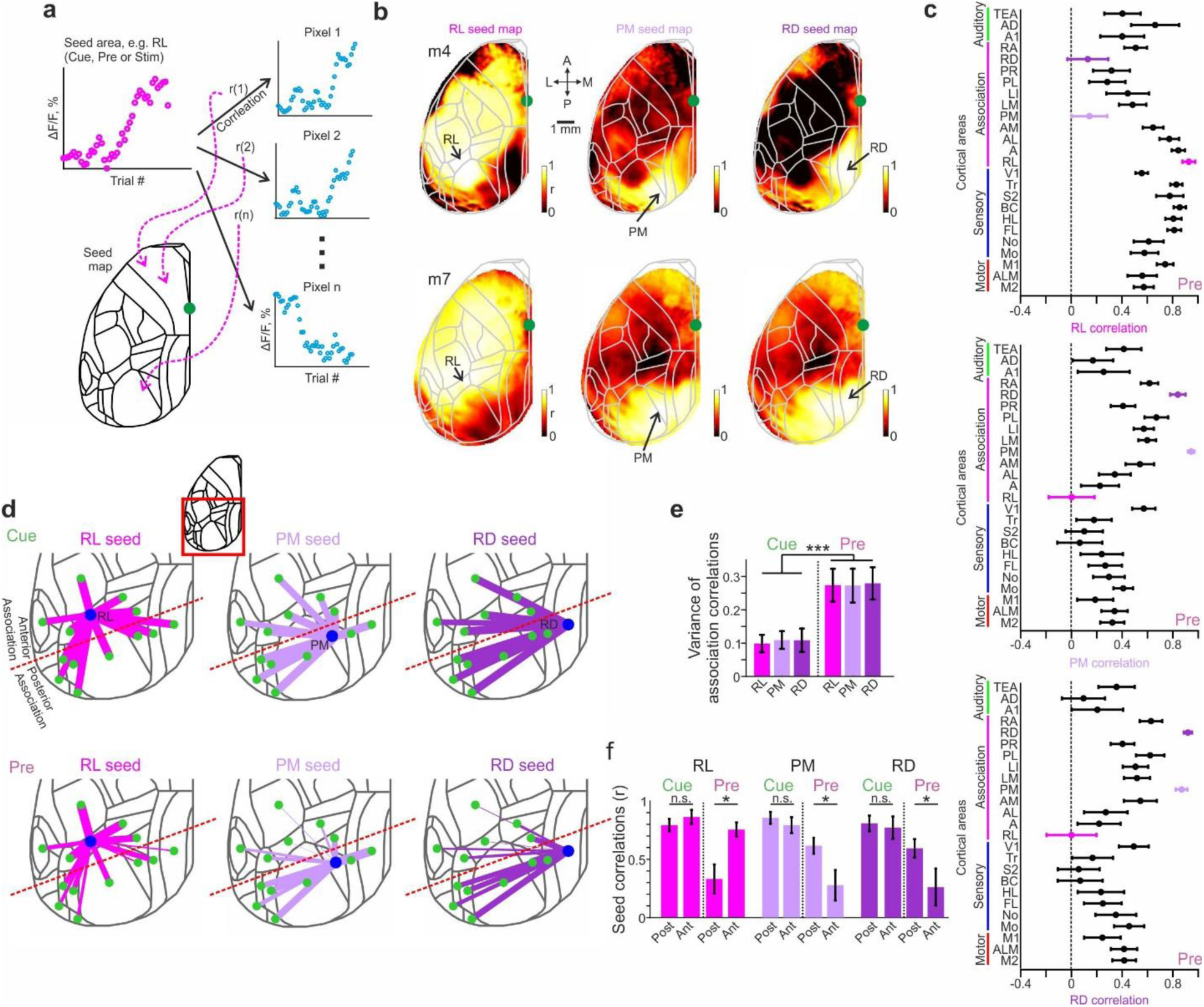
Functional reorganization of association cortex during learning. **a**, Schematic of calculating a seed correlation map. Each pixel in the map reflects the correlation coefficient (r) between the learning curve of a seed area (e.g. RL, PM or RD) and the learning-related ΔF/F signal changes of the respective pixel. Seed maps can be calculated separately for cue-, pre-, or stim-period. **b,** Seed maps for RL, PM and RD during pre-period in two example mice. Color denotes r-values. **c,** Correlation between the three seed areas and all the other areas during pre-period, averaged across all mice. Error bars are s.e.m. across mice. **d,** 2D overview of the inter-areal correlations across learning during cue-(top) and pre-period (bottom) between RL (left), PM (middle), and RD (right) and the surrounding cortical areas (also including A1, BC and V1). Line width is proportional to the average r-value across all mice. Red dashed line indicates separation into anterior and posterior association areas. **e,** Variance of inter-areal correlations within all association areas for each seed area in cue- and pre-period. Error bars are s.e.m. across mice. **f,** Seed area correlation values for the three areas during cue- and pre-period, averaged separately for anterior or posterior association areas. Error bars are s.e.m. across mice. *P < 0.05. ***P < 0.001. n.s. – not significant. Wilcoxon signed-rank test.

A closer look at the inter-areal correlations in the posterior part of cortex during cue- and pre-period revealed that all association areas are highly correlated during the cue-period but vary dramatically during the pre-period (**Fig. 6d**). For further quantification, we divided the association cortex into anterior (RL, A, AM and AL) and posterior (PM, RD, LM, LI, PL, PR and RA) areas (see dashed red line in Figure 6c). For all three seed areas the variance of correlation with other association areas was higher during the pre-period compared to the cue-period (**Fig. 6e**; p<0.001; Wilcoxon signed-rank test). Moreover, during the pre-period, unlike the cue-period, RL displayed significantly higher correlation with anterior compared to posterior association areas whereas PM and RD displayed the opposite effect (**Fig. 6f**; p<0.05 for the pre-period; p>0.05 for the cue-period; Wilcoxon signed-rank test; see a full correlation matrix for the learning-related ΔF/F changes in all 25 areas and for all 3 trial periods in **Supplementary Figure 8**). In summary, we find that the network of association areas is reorganized and spatially refined during learning by enhancing anterior association areas in a correlated manner while maintaining suppression in posterior association areas, specifically in the trial period before texture touch.

### Barrel cortex discriminates best between go and no-go textures

So far, we have mainly concentrated on go trials and on how large-scale cortical dynamics changes during learning. Most of the effects (e.g. widespread suppression followed by specific enhancement involving RL) related to the trial periods before the texture touches the whiskers and thus were also present for the no-go trials. But which areas can eventually develop discriminative power to distinguish between the two textures? Based on previous studies we first concentrated on the primary sensory area, i.e. BC^2,7,10,11^. The average time courses of trial-related ΔF/F signals in BC for go and no-go textures did not differ in the naïve mice (**Fig. 7a**). In the expert phase, in contrast, touch-evoked ΔF/F changes were generally enhanced with the response to the go-texture being substantially higher than for the nogo-texture (**Fig. 7a**). To calculate the discrimination power between go and no-go textures we computed receiver operating characteristic (ROC) curves of single trials^27,32^, with the area under the curve (AUC) relating to discrimination power. AUC values in BC showed a significant increase during the stim-period in the expert but not in the naïve phase (**Fig. 7b**). High AUC values in BC developed across learning during the stim-period (**Fig. 7c**). Finally, we calculated AUC values during the stim-period for all 25 areas in naïve and expert mice. Pooled across all mice, BC displayed the highest AUC values, followed by other areas mostly in motor, sensory and frontal association areas (**Fig. 7d**). Highest discrimination power in BC is further highlighted when calculating AUC values for each pixel to obtain an AUC map (**Fig. 7e**). Next, we further investigated the development of AUC in BC across learning (during the stim-period; **Fig. 6f**). In all mice, AUC values increase with learning and the learning threshold of each mouse was correlated with the inflection point of a sigmoidal fit to the AUC curve (**Fig. 7f,g**; r = 0.98; p<0.05). Thus, discrimination power in BC emerges at the same time when mice pass the learning threshold, indicating the tight link of the changes in cortical processing with improved task performance.

**Fig. 7.**
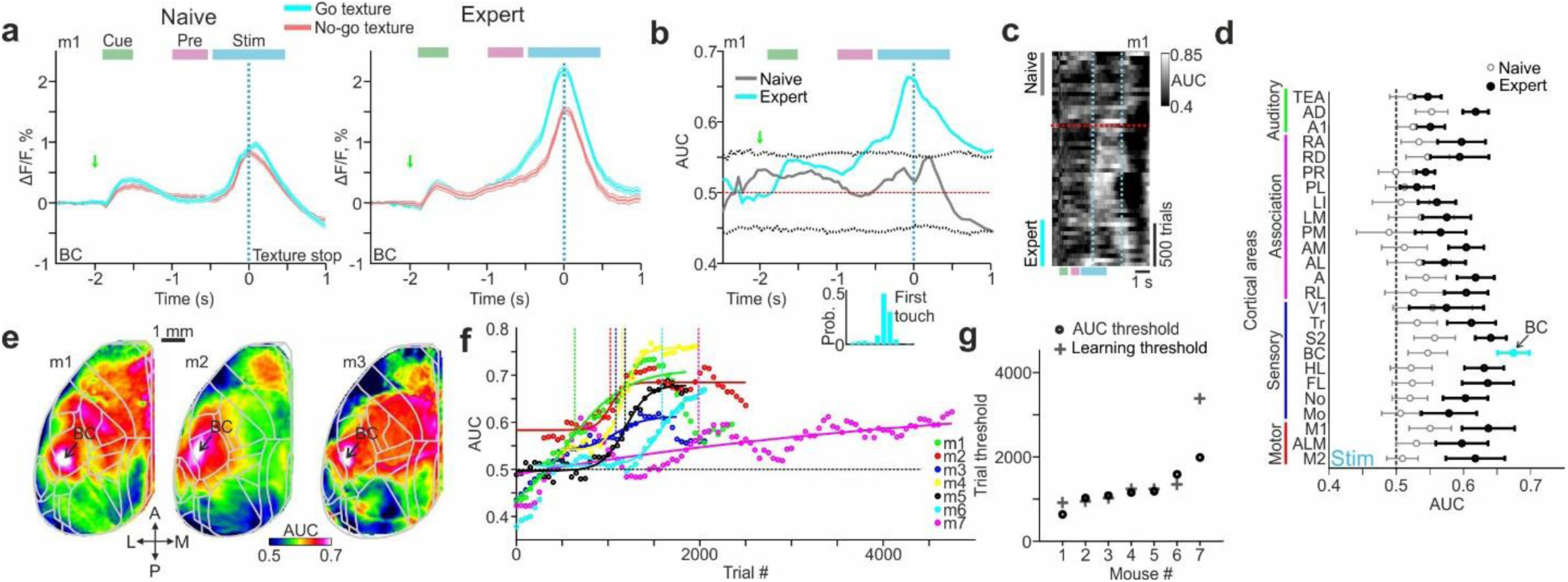
Emergence of discrimination power in barrel cortex during learning. **a**, BC activity in an example mouse for go (cyan) and no-go (red) trials in naïve (left) and expert (right) mouse. Error bars are s.e.m. across trials. **b,** ROC-AUC values for go vs. no-go trials as a function of time for naïve (gray) and expert (cyan) mice. Dashed gray lines indicate mean ± 2 s.d. of shuffled data. Same example as in a. Histogram below depicts the distribution of the first touch for this example. **c,** Heat map of trial-related AUC values across learning dimension (vertical axis) in the same example mouse. Dashed red line indicates learning threshold. Dashed cyan lines indicate texture-in period. **d,** Pooled AUC values during stim-period in all areas for expert (blue) and naïve (black) mice. Error bars are s.e.m. across mice. **e,** AUC maps during stim-period in three example expert mice. Color denotes AUC values. **f,** AUC values in BC during stim-period across learning for all mice. Threshold for each mouse is indicated with a vertical line at the inflection point of the sigmoid fit. **g,** Inflection points of AUC curves defining ‘AUC thresholds’. Learning thresholds are marked with gray plus signs.

## Discussion

We have identified learning-related changes in cortical activity covering a wide range in spatiotemporal space. First, changes were distributed across many cortical areas and they comprised suppression, enhancement, as well as sequential combinations thereof. Second, these changes were observed in the early trial periods before the texture-touch, indicating that an essential part of learning the animal needs to grasp the experimental setting and to understand the trial structure, within which the relevant stimulus for discrimination (here the texture touch) is embedded. Third, decreases in cortical activity occurred consistently several hundreds trials before the actual learning, suggesting that preparatory changes are required for the subsequent cortical adaptations that then presumably underlie the improvement of performance. The main pattern we observed is an early widespread suppression in association areas followed by an enhancement of a spatially more confined set of task-relevant areas (**Fig. 8**). Eventually, the emergence of a robust trial-related activation sequence from auditory cortex to RL to barrel cortex leads to the highest neural discrimination power in barrel cortex upon touch.

**Fig. 8.**
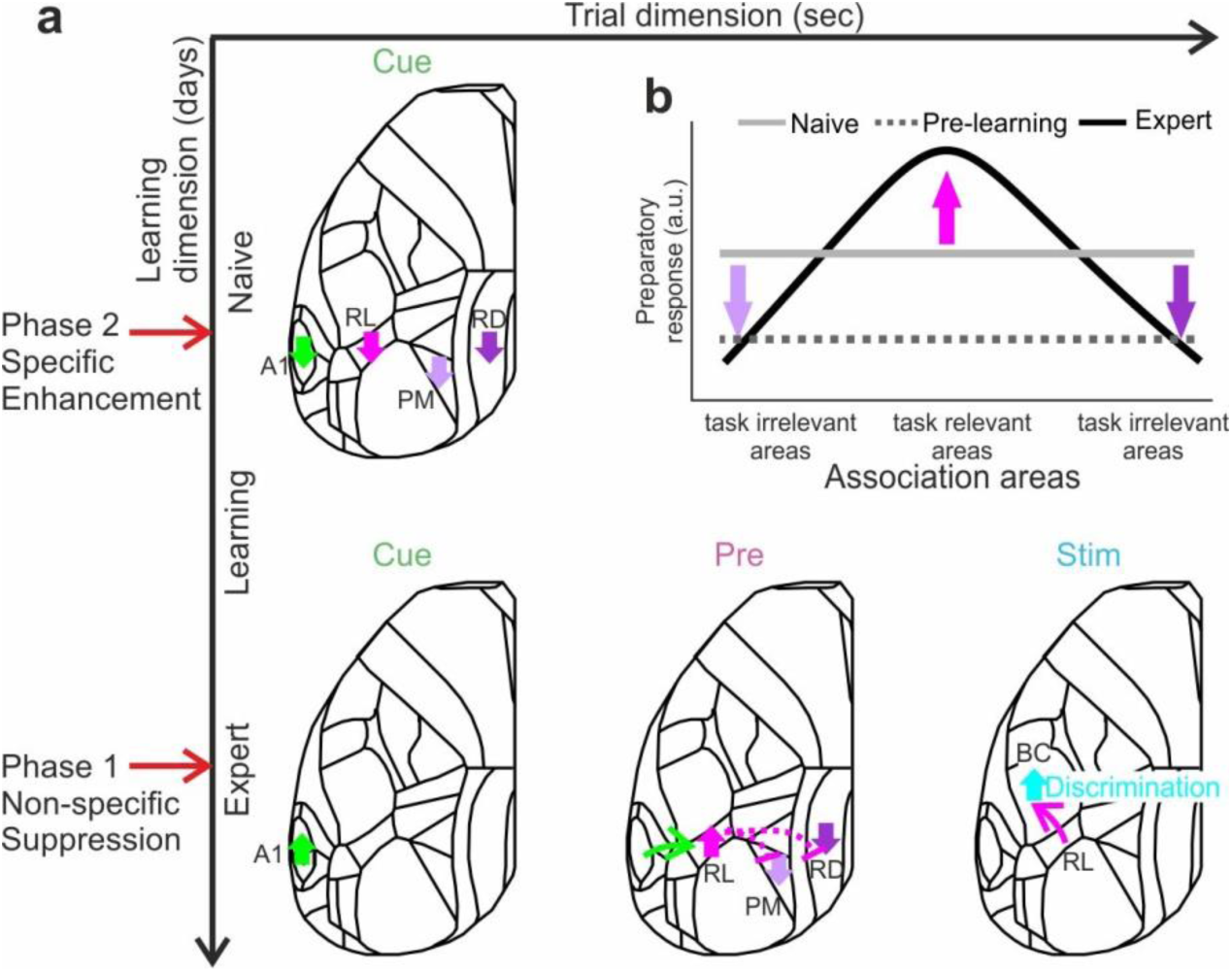
Learning starts with a general suppression phase followed by a specific enhancement phase. **a**, A schematic illustration of the main cortical changes within the two temporal scales: trial (x-axis) and learning (y-axis). Two phases occur across learning: before mice actually learn to discriminate between textures, association areas display non-specific suppression of activity in response to the cue (phase 1: non-specific suppression). Only later, as mice learn the task, a specific sequential pattern is enhanced: starting from A1 in response to the cue, then RL just before the texture touches the whiskers, and BC during texture touch (phase 2: specific enhancement). Association areas PM and RD specifically remain suppressed. **b**, Schematic diagram showing the distribution of responses across association areas, in preparation of the upcoming texture touch (pre period) for the naïve (gray) and expert (black).

The activation of RL (as part of PPC) before the texture touch may reflect predictive, anticipatory, expectation or attentional processes, as reported previously in PPC of primates (area LIP)^33,34^. Consistent with this notion, RL in mice displays predictive responses in the absence of the texture stimulus^35^ (see also **Supplementary Fig. 3**). In addition, anatomical projections from RL to BC^29,36^ imply top-down processing that may aid the preparation of adequate processing of the incoming relevant texture stimulus in BC and the association between the stimulus and a possible future reward. This is in line with our finding that the establishment of a robust temporal sequence with RL bridging the stimulus-cue to the task-relevant texture stimulus follows a similar time course as the divergence of BC signals for Hit and CR trials. The emergence of discrimination power in BC also went hand in hand with increases in body movements, apparently in expectation of and preparation for the upcoming reward. These extensive movements result in widespread and large cortical activity, including forelimb cortex and motor areas, which makes it difficult to separate task-related stimulus processing from behavior-related activation patterns, as they increasingly mix in the later trial phase after the touch has happened. Further experiments will be required to dissect the cortical signal flow related to the conversion of touch information into preparatory and executive motor signals.

Regarding the temporal dimension of learning that spans several days, we found a general suppression of several association areas, e.g. PM and RD, as a particularly salient event (**Fig. 8**). This early suppression was obvious around 500 trials before a mouse learns to discriminate between the textures, during a time period when the mean performance is still low (d’ = 0.32 ± 0.31 mean ± s.e.m.; most mice still lick for both textures). Because these suppression events occurred consistently before onset of learning, we could even predict when the mouse is likely to reach learning threshold based on this dynamic feature. In our interpretation, suppression during the cue-period may indicate a general attentive state that is a prerequisite for learning. It is likely that such increased attention is also apparent in the animal’s behavior and we indeed observed a decrease in obsessive licking during the reward period, which occurred about at the time when the suppression was observed. In auditory cortex we found a combination of early suppression followed by later enhancement, indicating that mice may use the stimulus cue information in order to prepare for the upcoming trial. There was no significant difference in the A1 response to the stimulus cue between expert and naïve mice but a clear reduction during the steepest part of the learning curve (Fig. 3e). This result points to a pronounced reorganization during this training phase and also emphasizes the dynamic process that is induced during learning in order to feed anticipatory signals into the task-relevant area.

Interestingly, we found dissociation of signaling between anterior and posterior sets of association areas, which may reflect different roles of these areas in neural processing. Anatomical evidence suggests that RL uni-directionally projects to PM^36^, which possibly could exert inhibitory control via interneurons. PM and RD are strongly bi-directionally connected, but less so with anterior association areas^29,36^, thus substantiating our functional findings. What could be the reason for the pronounced and persistent suppression of PM and RD? PM has been studied mostly in the context of visual tasks highlighting its role in spatial processing and navigation^37–39^. RD is also connected with hippocampal regions and has been shown to convey top-down effects in a visual discrimination task^4^ and has been linked to spatial navigation and memory^40–42^. Therefore, it may be that under our experimental conditions, where spatial navigation is not relevant, this network is actively suppressed. Future studies may investigate whether in tasks distinct from ours, where spatial aspects are important, posterior association areas may show enhanced activity while anterior association areas including RL may be suppressed. Alternatively, RL could be the association area for processing tactile information whereas PM serves as association area for visual information and is not needed in our task. In summary, our results highlight the distributed functional reorganization that cortical dynamics undergoes during learning, progressing in two distinct major phases that first reflect the transition into a ready-for-learning modus and second establish the specific cortical flow pattern that is needed to solve the task.

## ACKNOWLEDGEMENTS

This work was supported by grants from the Swiss National Science Foundation (SNSF) (31003A- 149858; F.H.), the European Research Council (ERC Advanced Grant BRAINCOMPATH, project 670757; F.H.), an Edmond and Lily Safra Center for Brain Sciences (ELSC) postdoctoral fellowship (A.G.), an EMBO long-term postdoctoral fellowship (ALTF_1077-2014; A.G.), and a Marie-Curie Individual Fellowship (659719-AG-GF; A.G.).

## AUTHOR CONTRIBUTIONS

A.G. and F.H. designed the experiments. A.G. conducted the experiments. A.G. and F.H. performed data analysis. A.G. and F.H. wrote the manuscript.

## DATA AVAILABILITY

The data and code that support the findings of this study are available from the corresponding author upon reasonable request.

## METHODS

### Animals and surgical procedures

Methods were carried out according to the guidelines of the Veterinary Office of Switzerland and following approval by the Cantonal Veterinary Office in Zurich. A total of 7 adult male mice (1-4 months old) were used in this study. These mice were triple transgenic Rasgrf2-2A-dCre;CamK2a-tTA;TITL-GCaMP6f animals, expressing GCaMP6f in excitatory neocortical layer 2/3 neurons^11^. To generate triple transgenic animals, double transgenic mice carrying CamK2a-Tta ^43^ and TITL-GCaMP6f ^44^ were crossed with a Rasgrf2-2A- dCre line (^45^; individual lines are available from The Jackson Laboratory as JAX# 016198, JAX#024103, and JAX# 022864, respectively). The Rasgrf2-2A-dCre;CamK2a-tTA;TITL- GCaMP6f line contains a tet-off system, by which transgene expression can be suppressed upon doxycycline treatment^46,47^. However, doxycycline treatment is not necessary in these animals, since the Rasgrf2-2A-dCre line holds an inducible system of its own, given that the destabilized Cre (dCre) expressed under the control of the Rasgrf2-2A promoter needs to be stabilized by trimethoprim (TMP) to be fully functional. TMP (Sigma T7883) was reconstituted in Dimethyl sulfoxide (DMSO, Sigma 34869) at a saturation level of 100 mg/ml, freshly prepared for each experiment. For TMP induction, mice were given a single intraperitoneal injection (150 µg TMP/g body weight; 29g needle), diluted in 0.9% saline solution.

We used an intact skull preparation^48^ for chronic wide-field calcium imaging of neocortical activity which we previously described^11^. Mice were anesthetized with 2% isoflurane (in pure O_2_) and body temperature was maintained at 37°C. We applied local analgesia (lidocaine 1%), exposed and cleaned the skull, and removed some muscles to access the entire dorsal surface of the left hemisphere (Figure 2A; ∼6 x 8 mm^2^; from ∼3 mm anterior to bregma to ∼1 mm posterior to lambda; from the midline to at least 5 mm laterally). We built a wall around the hemisphere with adhesive material (iBond; UV-cured) and dental cement “worms” (Charisma). Then, we applied transparent dental cement homogenously over the imaging field (Tetric EvoFlow T1). Finally, a metal post for head fixation was glued on the back of the right hemisphere. This minimally invasive preparation enabled high-quality chronic imaging with high success rate.

### Texture discrimination task

Mice were trained on a go/no-go discrimination task (Fig. 1a). The behavioral setup has been described previously^27^. Each trial started with an auditory cue (stimulus cue; 2 beeps at 2 kHz, 100-ms duration with 50-ms interval), signaling the approach of either two types of sandpapers (grit size P100: rough texture; P1200: smooth texture) to the mouse’s whiskers as ‘go’ or ‘no-go’ textures (Fig. 1a; pseudo-randomly presented with no more than 3 repetitions). The texture stayed in touch with the whiskers for 2 seconds, and then it was moved out after which an additional auditory cue (response cue; 4 beeps at 4 kHz, 50-ms duration with 25-ms interval) signaled the start of a 2 second response period. A water reward was given to the mouse for licking for the go texture only after the response cue (‘hit’). Punishment with white noise was given for licking for the no-go texture (‘false alarms’; FA). Licking before the response cue was not rewarded or punished. Reward and punishment were omitted when mice withheld licking for the no-go (‘correct-rejections’, CR) or go (‘Misses’) textures. The licking detector remained in a fixed and reachable position throughout the entire trial. Note that the auditory tones merely served as cues defining the temporal trial structure, but had no predictive power with respect to go or no-go condition. The first auditory tone signaled the trial-start and thus predicted the upcoming arrival of the texture as the task-relevant stimulus, whereas the second auditory tone indicated the availability of a water reward in the go trials. Licking before the response cue was allowed and did not lead to punishment or early reward.

#### Training and performance

Five mice were trained to lick for the P100 texture (mice #1-4 and 7) and 2 mice were trained to lick for the P1200 texture (mice #5 and 6). Mice were first handled and accustomed to head fixation before starting water scheduling. Before imaging began mice were conditioned to lick for reward after the presentation of the texture. Imaging began only after mice reliably licked for the response cue (typically after the 1^st^ day; 200-400 trials). On the first day of imaging, mice were presented with the ‘go’ texture and after 50 trials the ‘no-go’ texture was gradually introduced (starting from 10% and increasing by 10% approximately every 50 trials;^49^) until reaching 50% probability for the no-go texture by the end of the day. During the 2^nd^ day, most mice continuously licked for both textures (supplementary Fig. 2). Thus after around 100 trials, we increased no-go probability to 80% and waited for mice to perform three continuous CR trials before returning to 50% probability. This was done for several times until mice increased their performance, specifically withheld licking for the no-go texture. In mice that still continued to lick for both textures we additionally repeated the wrong response until a correct response. In all mice, a 50% protocol was presented with no repetitions as soon as they reached expert level (d’>1.5). 6 out of the 7 mice learned the task within 3-4 days after around a thousand trials (Fig. 1d; supplementary Fig. 2). Mouse #7 learned the task within 10 days. An effort was made to maintain a constant position of the texture and cameras across imaging days in order to maintain similar stimulation and imaging parameters.

### Wide-field calcium imaging

We used a wide-field approach to image large parts of the dorsal cortex while mice learned to perform the task^11^. A sensitive CMOS camera (Hamamatsu Orca Flash 4.0) was mounted on top of a dual objective setup. Two objectives (Navitar; top objective: D-5095, 50 mm f0.95; bottom objective inverted: D-2595, 25 mm f0.95) were interfaced with a dichroic (510 nm; AHF; Beamsplitter T510LPXRXT) filter cube (Thorlabs). This combination allowed a ∼9 mm field-of-view, covering most of the dorsal cortex of the hemisphere contralateral to texture presentation. Blue LED light (Thorlabs; M470L3) was guided through an excitation filter (480/40 nm BrightLine HC), a diffuser, collimated, reflected from the dichroic mirror, and focused through the bottom objective approximately 100 µm below the blood vessels. Green light emitted from the preparation passed through both objectives and an emission filter (514/30 nm BrightLine HC) before reaching the camera. The total power of blue light on the preparation was <5 mW, i.e., <0.1 mW/mm^2^. At this illumination power we did not observe any photo-bleaching. Data was collected with a temporal resolution of 20 Hz and a spatial resolution of 512×512. On each imaging day a green reflectance image was taken as reference to enable registration across different imaging days using the blood vessel pattern (fiber-coupled LED illuminated from the side; Thorlabs).

#### Mapping and area selection

Each mouse underwent a mapping session under anesthesia (1% isoflurane), in which we presented five different sensory stimuli (contra-lateral side): a moving bar stimulating multiple whiskers, the forelimb paw, or the hindlimb paw (20 Hz for 2 s); visual stimulation with a blue LED in front of the eye (100 ms duration; approximately zero elevation and azimuth); and white noise auditory stimulation (2 s. duration). The averaged evoked maps clearly showed activation patches in the expected areas (Fig. 1c; supplementary Fig. 1a). Next, we registered each imaging day to the mapping day using skull coordinates from the green images. Finally, we registered each mouse onto a 2D top view mouse atlas using both functional patches from the mapping and skull coordinates (supplementary Fig. 1). Within the atlas borders, we defined 25 areas of interest, with some manual modifications within these borders to fit the functional activity for each mouse. Motor cortex areas were defined based on stereotaxic coordinates and functional patches for each mouse (see below). Thus all mice had similar regions of interest that were comparable within and across mice.

We grouped these 25 areas into auditory (green; Au), association (pink; Asc), sensory (blue; somatosensory along with primary visual cortex; SV), and motor (red; M) areas (Fig. 1d and supplementary Fig. 1b). Auditory areas: Primary auditory (A1), Auditory dorsal (AD) and Temporal association area (TEA). Senosory areas: Somatosensory mouth (Mo), Somatosensory nose (No), Somtosensory hindlimb (HL), Somtosensory forelimb (FL), Barrel cortex (BC; primary somatosensory whisker); Secondary somatosensory whisker (S2), Somtosensory trunk (Tr) and Primary visual cortex (V1). Motor areas: whisker-related primary motor cortex (M1; 1.5 anterior and 1 mm lateral from bregma, corresponding to the whisker evoked activation patch in M1 from the mapping session), anterior lateral motor cortex (ALM; 2.5 anterior and 1.5 mm lateral from bregma^50^) and secondary motor cortex (M2; 1.5 anterior and 0.5 mm lateral from bregma corresponding;^11^). Association cortex: Rostrolateral (RL), Anterior (A), Anterior lateral (AL), Anterior medial (AM), Posterior medial (PM), Lateral medial (LM), Lateral intermediate (LI), Posterior lateral (PL), Post-rhinal (PR), Retrosplenial dorsal (RD) and Retrosplenial angular (RA). We note that our definition of association cortex is broad and may include or exclude areas that are not necessarily classical association areas. In addition, we further divided association areas into anterior (RL, A, AM and AL) and posterior (PM, L, LI, PL, PR, RD and RA) association cortex (dashed red line in Fig. 5d).

#### Control experiments

In control experiments, we excluded confounding effects of autofluorescence or non-calcium-related intrinsic signals, by exciting the wide-field preparation with green light, showing no positive responses during cue-, pre-, and stim-period (**Supplementary Fig. 9**; For additional controls for non-calcium related optical signals see^11^). Therefore, in the experiments presented in this study non-calcium-related intrinsic signals have no major influence on the GCaMP6f signals, especially in the cue and pre periods. To control for possible changes in responses across several days that are not necessarily related to learning, we evaluated the stability of areal activity in expert mice imaged across 5 consecutive days. Responses in BC (during stim-period), RL (during pre-period), and A1 (during cue-period) across 5 days were relatively flat (n = 4 mice). In addition, trial-shuffled data across learning eliminated these changes in responses and resulted in a relatively flat change in response (10^3^ iterations). Taken together, changes in activity across a learning period of several days is more likely to be learning related rather than day-to-day fluctuations in activity.

### Whisker and body tracking

In addition to wide-field imaging, we tracked movements of the whiskers and the body of the mouse during the task (Fig. 1a). The mouse was illuminated with a 940-nm infra-red LED. Whiskers were imaged at 50 Hz (500×500 pixels) using a high-speed CMOS camera (A504k; Basler), from which we calculated time course of whisking envelope and the time of first touch (see below). An additional camera monitored the movements of the mouse at 30 Hz (The imaging source; DMK 22BUC03; 720×480 pixels). We used movements of both forelimbs and the head/neck region to assess body movements, to reliably detect large movements (Fig. 1a; see *Data Analysis* below).

### Data analysis

Data analysis was performed using Matlab software (Mathworks). All mice were continuously imaged during learning (5-11 days). Wide-field fluorescence images were sampled down to 256×256 pixels and pixels outside the imaging area were discarded. Each pixel and each trial were normalized to baseline several frames before the stimulus cue (frame 0 division). In this study, we grouped trials based on the texture type, i.e. go or no-go texture (see *Calculating learning curves* below). We define three time period within the trial structure: cue (−1.9 to −1.6 relative to texture stop), pre (−1 to 0.5 s relative to texture stop) and stim (−0.5 to 0.5 relative to texture stop; Fig. 1d). Naïve and expert mice are defined as the first and last 500 trials respectively.

#### Calculating body movements

We used a body camera to detect general movements of the mouse (30 Hz frame rate; supplementary Fig. 1a). For each imaging day, we first outlined the forelimbs and the neck areas (one area of interest for each), which were reliable areas to detect general movements. Next, we calculated the body movement (1 minus frame-to-frame correlation) within these areas as a function of time for each trial. Thresholding at 3 s.d. (across time frames before stimulus cue) above baseline (defined as the 5^th^ percentile) resulted in a binary movement vector (either ‘moving’ or ‘quiet’) for each trial^11^.

#### Whisker tracking and first-touch analysis

The average whisker angle across all imaged whiskers was measured using automated whisker tracking software^51^. The mean whisker envelope was calculated as the difference between maximum and minimum whisker angles along a sliding window equal to the imaging frame duration (50 ms;^11,27^). Whisker envelope was normalized just before the auditory cue similar to widefield data (Frame zero). In addition, we manually detected the first frame, in which any whisker touched the upcoming texture, using the movies from the whisker cameras (LabVIEW custom program). The first touch occurred on average 0.33 and 0.34 s before the texture stopped for naïve and expert mice respectively. Time of first touch did not differ between expert and naïve mice (*P* > 0.05; Mann-Whitney U-test; n=7 mice). We note that the pre period from −1 to −0.5 relative to texture stop mostly (but not exclusively) does not contain the first touch.

#### Calculation of curves across learning

Trials were binned (n=50 trials with no overlap) across learning and the performance (defined as d’ = *Z*(Hit/(Hit+Miss)) – *Z*(FA/(FA+CR)) where *Z* denotes the inverse of the cumulative distribution function) was calculated for each bin. Next, each behavioral learning curve was fitted with a sigmoid function

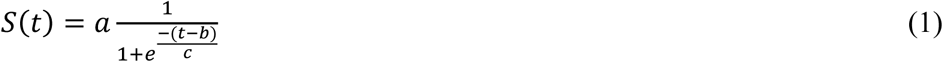

Where *a* denotes the amplitude, *b* the time point (in trial numbers) of the inflection point, and *c* the steepness of the sigmoid. A d’=1.5 was defined as the threshold and mice were ordered based on the trial number in which they crossed threshold (i.e. learning threshold; Fig. 1g). Different threshold maintained the order of the mice based on their learning threshold (see Fig. 1f).

To compare the behavioral learning curve with other behavioral parameters and neuronal activity, we similarly grouped trials and separated them based on the texture type, i.e. hit and miss trials were grouped into the go texture; CR and FA trials were grouped into the no-go texture. Our main focus in this study was on the go texture (presented in figures 1-6). Therefore, stimulus identity was kept similar across learning. Results were maintained when considering only the no-go texture. Only in figure 7 we compare between go and no-go textures to calculate discrimination power. Next, we can present a behavioral parameter (i.e. body movement, whisking envelope or licking probability) or cortical areas (averaged over pixels) in 2 dimensional temporal spaces where the x-axis is the trial temporal structure (i.e. trial dimension) and the y-axis is the learning profile across trials and days (i.e. learning dimension; for examples see Fig. 2a, e, i; Fig. 3c top). From this 2D temporal space we could average across trials of the learning dimension, e.g. during naïve and expert states (for example see Fig. 2b, f, j; Fig. 3c middle). Alternatively, we can average across time frames within the trial dimension, to obtain a response curve across learning for a specific time period (i.e. cue, pre or stim period; additionally smoothed with a Gaussian kernel (2σ=9) and fitted with a sigmoid function; for example see Fig. 2c, g, k; Fig 3c bottom). Thus we are able to obtain a curve across learning for a specific area or behavioral parameter which are comparable to the behavioral learning curve of the mouse. The sigmoid fits of the response curves from different cortical areas were normalized between 0 and 1 in order to compare between response curves of different areas. This was done mainly because of the different activation ranges across learning for each area. Non-normalized learning curves are presented in supplementary Figure 6. In an additional analysis we also fitted each response curve for all areas and time periods with a double sigmoid fit in order to fit both the initial suppression and the later enhancement that was present in some curves (e.g. Fig. 4d; Supplementary Fig. 8):

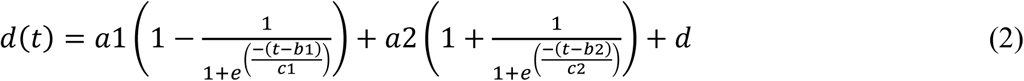

with *a1* and *a2* as amplitudes, *b1* and *b2* as inflection points (in trial numbers), and *c1* and *c2* as steepness parameters of the descending and ascending sigmoid. respectively. *d* is baseline parameter, which was set to the minimum value of a curve. Thus, for each area we could quantify the amount (amplitude) and timing (latency) of both suppression and enhancement during each time period relative to the learning threshold. Finally, to quantify the enhancement suppression ratio we calculated the modulation index (MI) as

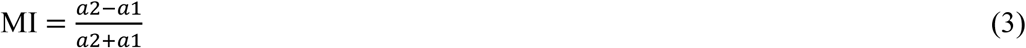

ranging from −1 and 1, with positive values indicating more enhancement, negative values indicating more suppression, and near zero values indicating similar amounts of suppression and enhancement.

#### Calculating learning maps and seep maps

To study the relationship between the behavioral learning curve and the learning curves of all pixels we calculated a learning map (Fig. 5). This was done by calculating the correlation coefficient (r) between the behavioral learning curve of the mouse and the learning-related ΔF/F changes of each pixel (Fig. 5a). This can be done for a specific time period (i.e. cue, stim or stim periods; Fig. 5a, b) or for each time frame (Fig. 5c). To calculate the relationship between the learning-related ΔF/F changes of a specific area (i.e. seed) and the learning-related ΔF/F changes of all pixels we calculated a seed correlation map (Fig. 6). This was done similarly to the learning map by only substituting the behavioral learning curve with the learning-related ΔF/F changes of the desired area (defined as the seed area; Fig. 6a). We chose seed areas to be RL, PM and RD which were of the most interest from previous analysis and best represent the main trends of neuronal changes during learning. A full correlation matrix between all learning curves is presented in Supplementary Fig. 8.

#### Discrimination power between go and no-go texture

To measure how well could neuronal populations discriminate between go and no-go textures, we calculated a receiver operating characteristics (ROC) curve and calculated its area under the curve (AUC). This can be done for each pixel (Fig. 7e), each area (Fig. 7d), each time frame (Fig. 7b), and across learning (Fig. 7f). Our main focus is during the stim period where texture touched the whiskers. To calculate significance, we calculated the sample distribution by trial shuffling between go and no-go textures (n=100 iterations). Exceeding mean±2std of the sample distribution is defined as significant (Fig. 7b).

#### Statistical analysis

In general, non-parametric two-tailed statistical tests were used, Mann-Whitney U-test to compare between two medians from two populations or the Wilcoxon signed rank test to compare a population’s median to zero (or between two paired populations). Multiple group correction was used when comparing between more than two groups.

