## Supplementary material for "Spatiotemporal refinement of signal flow through association cortex during learning": 9 Supplementary Figures

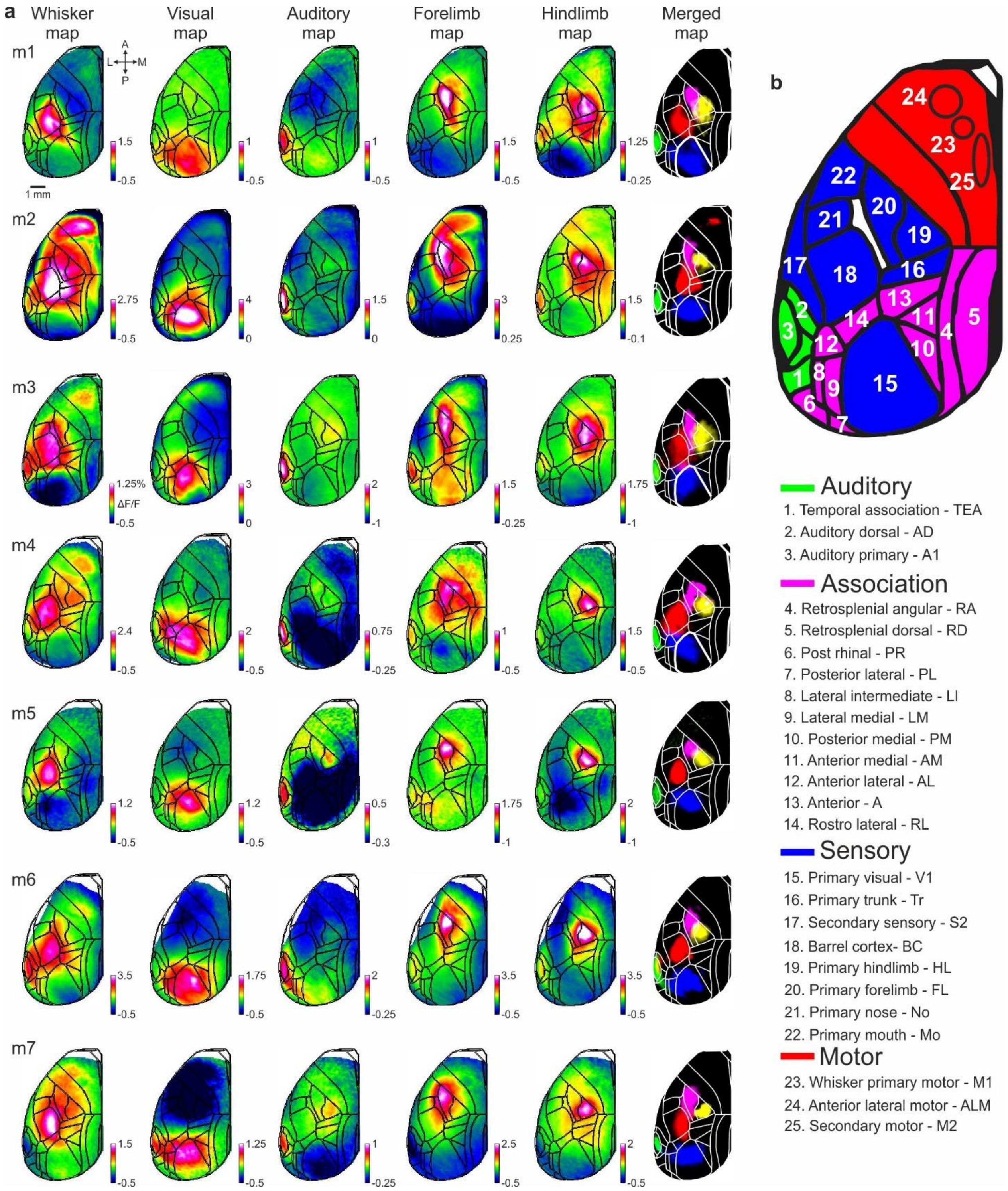

**Supplementary Figure 1 │ Functional mapping and area definitions. a,** A schematic illustration of the behavior and imaging setup. Additional cameras and detectors monitored body movements, whisking and licking of the mouse throughout the trial. **b,** Each mouse underwent a mapping session including five stimulus modalities: whisker, visual, auditory, forelimb and hindlimb (Methods). Mean activation maps are shown for each stimulus type for two example mice. Color denotes normalized fluorescence (ΔF/F). Furthermore, maps were registered onto a top 2D view of the mouse atlas (black lines). **c,** Full names and abbreviations of all the 25 areas used in this study. Areas were divided into auditory (green; Au), association (pink; Asc), sensory (blue; somatosensory and primary visual cortex; SV) and motor (M; red) cortices.

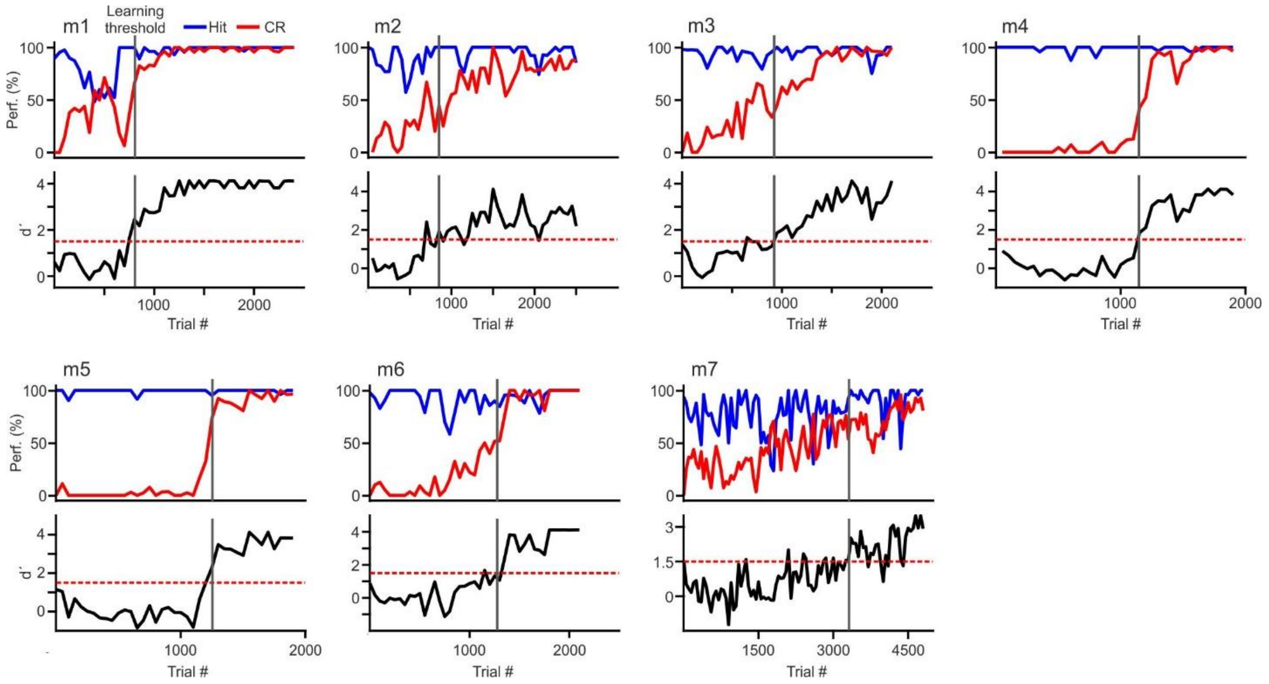


**Supplementary Figure 2 │ Behavioral performance and learning curves.** Behavioral performance for all seven mice, plotting the time course of Hit and CR rates in percent (top) and d' (bottom) as a function of trial number. The learning threshold when mice reached d' = 1.5 is indicated by vertical gray lines.

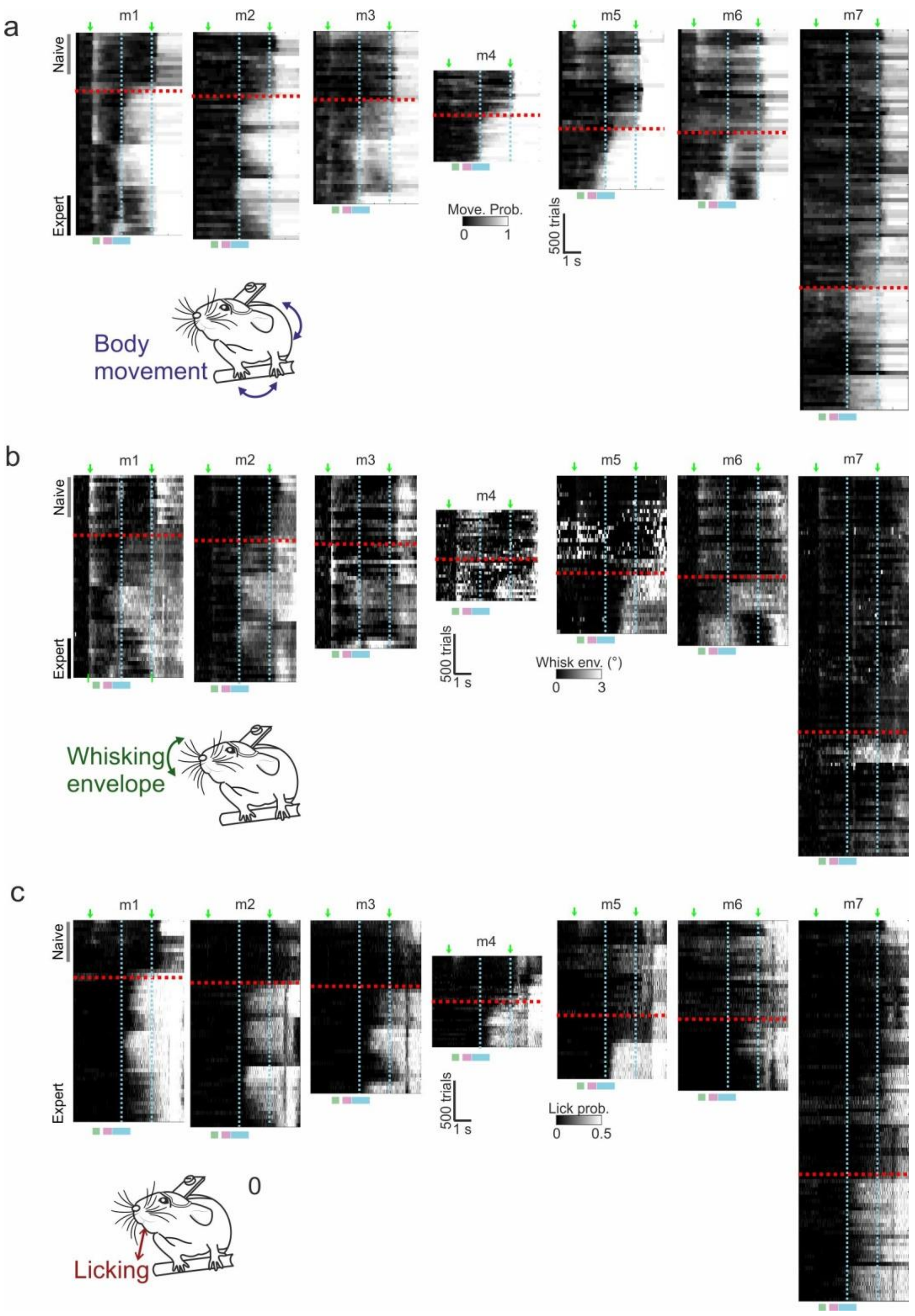


**Supplementary Figure 3 │ Behavioral changes during learning. a,** Body movements quantified as movement probability throughout the trial time across learning for all 7mice (50-trial bins; similar to Fig. 2a). Dashed red line indicates learning threshold. Heat maps are vertically aligned to this learning threshold. Dashed cyan lines indicate the texture-in period. Green arrows mark stimulus and response cues. **b,** Equivalent heat maps as in a but for the trial-related whisking envelope dynamics. **c,** Equivalent heat maps as in a but for the trial-related licking probability.

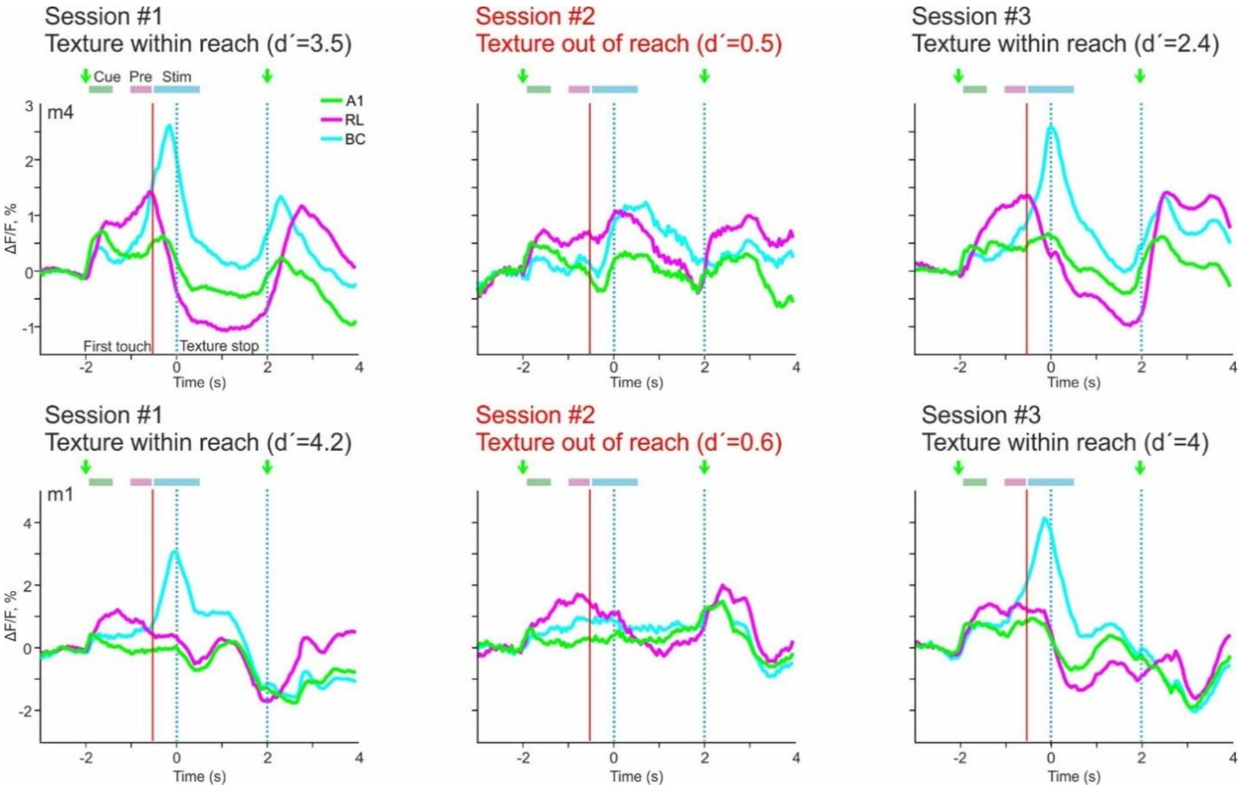


**Supplementary Figure 4 │Cortical activation responses in expert mice with the texture out of reach.** In two mice that reached expert level we performed wide-field calcium imaging when the texture was placed just out of reach of the whiskers. Here we plot the cortical signals in go trials for A1 (green traces), RL (magenta), and BC (cyan), comparing a session with texture out-of-reach (session #2) with the previous and the subsequent session, when the texture was within normal reach (sessions #1 and #3). Note in particular the strong activation response in RL during the pre-period, which persisted even when the texture was out-of-reach. In contrast, the texture-touch related activation observed in BC was abolished (as well as the subsequent suppression of RL and A1 during touch). The activations at the start of texture removal were also reduced although interestingly a response to the stimulus-cue only remained in RL in session #2.

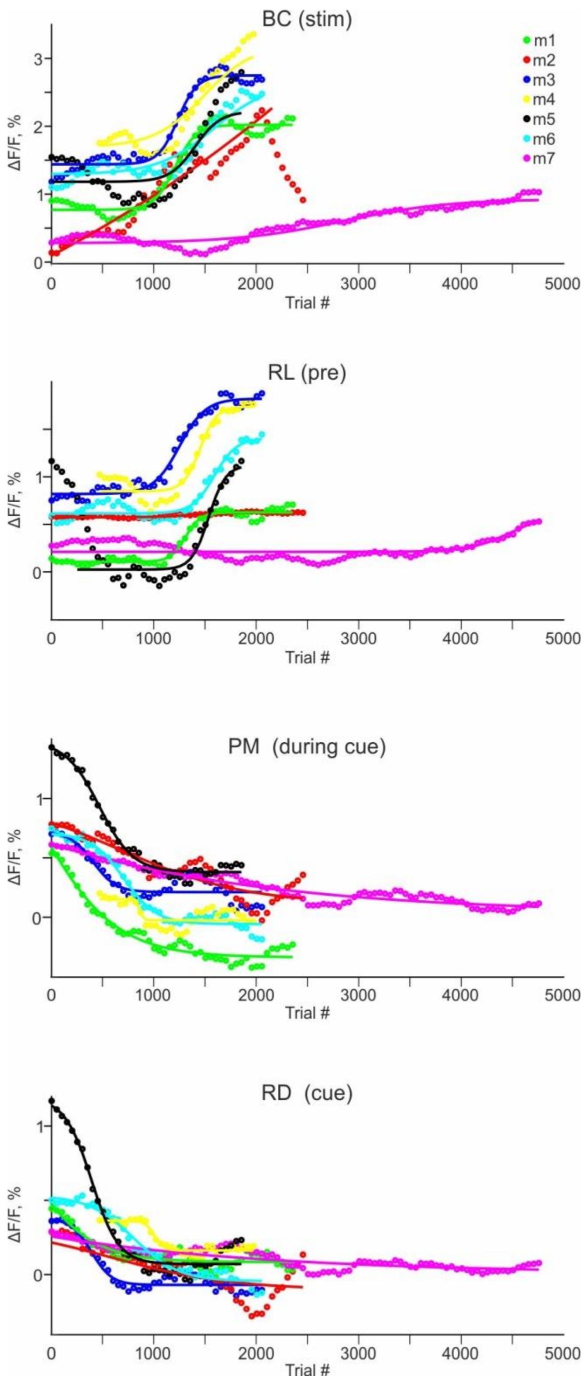


**Supplementary Figure 5 | Examples of non-normalized learning-related activity changes.** Learning-related changes in mean DF/F signals for four selected areas for each mouse (different colors). From top to bottom: BC during stim-period, RL during pre-period, PM during cue-period and RD during cue-period. Data were fitted with sigmoidal curves.

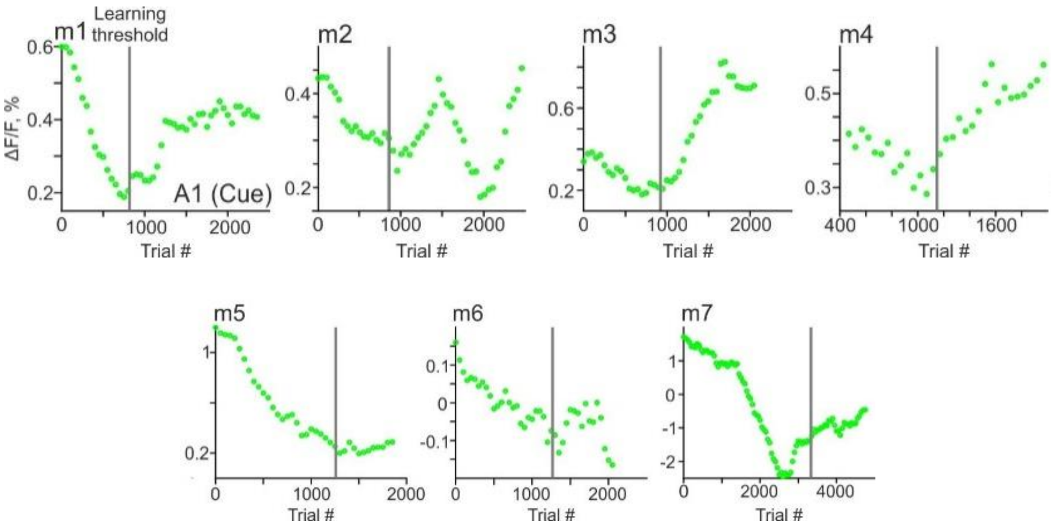


**Supplementary Figure 6 │ Variability of learning-related changes in A1.** Time course of learning-related changes in ΔF/F signals in A1 during the cue-period for all 7 mice. Learning thresholds are indicated with vertical gray lines. Note that suppression consistently occurs before the learning threshold while enhancement thereafter.

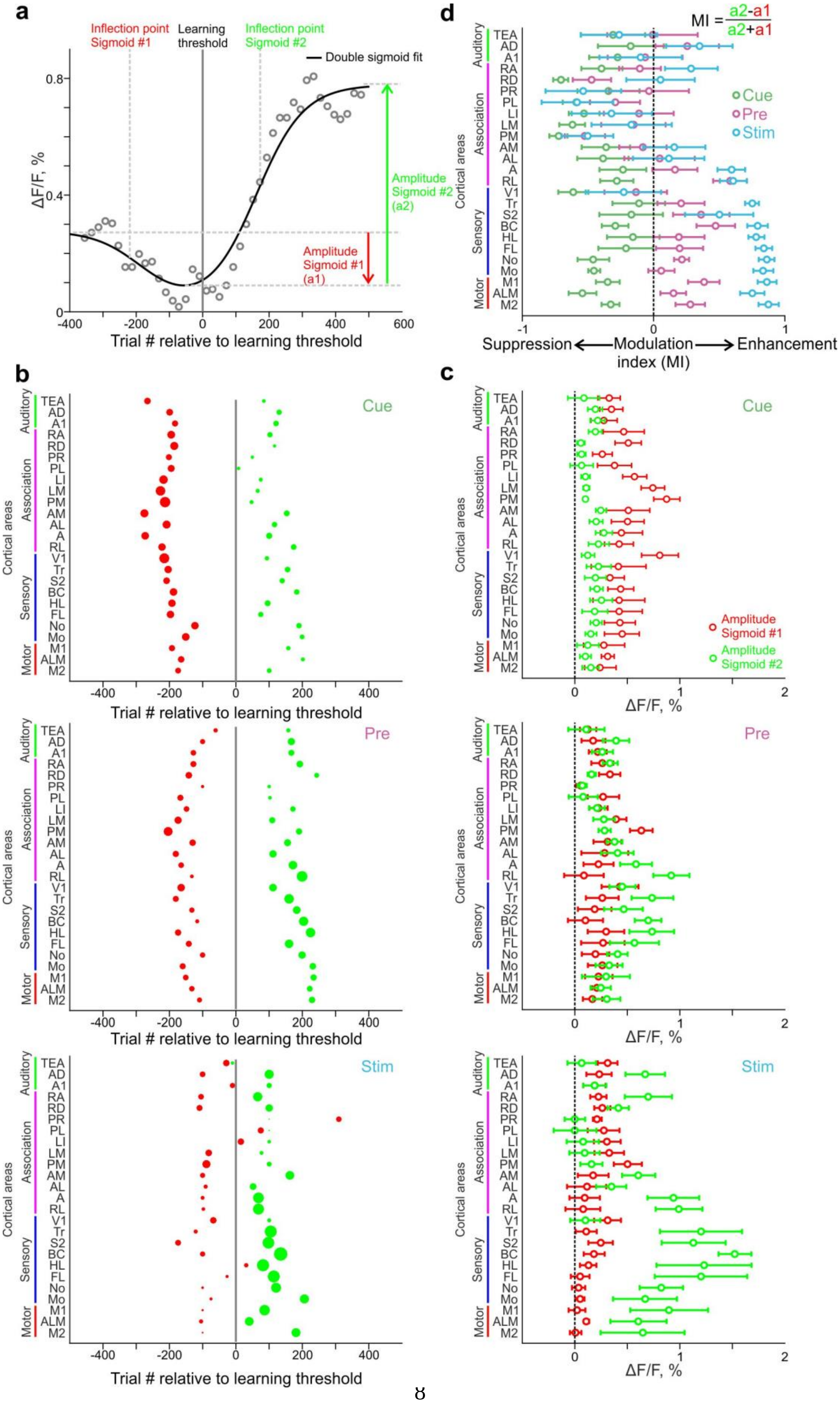
a
Inflection point Sigmoid #1
Learning threshold
Inflection point Sigmoid #2
Double sigmoid fit
ΔF/F, %
Amplitude Sigmoid #2 (a2)
Amplitude Sigmoid #1 (a1)
Trial # relative to learning threshold
b
Cue
Pre
Stim
Cortical areas
Auditory
Association
Sensory
Motor
TEA
AD
A1
RA
RD
PR
PL
LI
LM
PM
AM
AL
A
RL
V1
Tr
S2
BC
HL
FL
No
Mo
M1
ALM
M2
c
Amplitude Sigmoid #1
Amplitude Sigmoid #2
ΔF/F, %
d
MI = (a2-a1)/(a2+a1)
Cue
Pre
Stim
Suppression
Modulation index (MI)
Enhancement

**Supplementary Figure 7 │ Quantification of suppression and enhancement in the pre-learning and learning phases for all areas and trial periods. a.** An example learning curve from a given area (gray points) along with a double-sigmoid fit (black curve) comprised of a decreasing and an increasing sigmoid function (red and green, respectively). Four parameters are depicted: the amplitudes and the inflection points of the sigmoids. **b.** Grand average of amplitudes and latencies for the decreasing (red) and increasing (green) sigmoid for each area during the cue-, pre-, and stim-periods. Inflection points are arranged relative to the learning thresholds with the circle diameter proportional to the sigmoid amplitude. **c.** Amplitudes of each sigmoid for all areas in the cue-, pre- and stim-periods. Error bars are s.e.m across mice. **d.** Modulation index to quantify enhancement/suppression ratio for each area and each trial period. Positive values indicate predominant enhancement whereas negative values relate to predominant suppression. Error bars as in c.

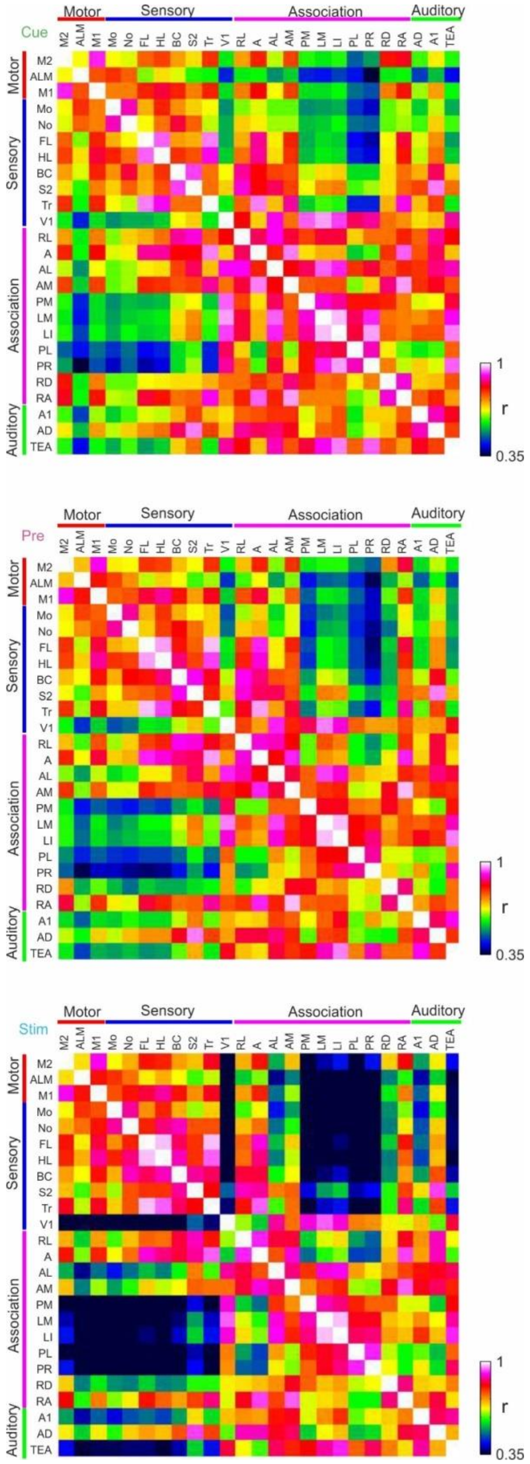


**Supplementary Figure 8 │ Correlation matrices between learning curves of different areas.** Full correlation matrix for all 25 areas, showing the pair-wise correlation coefficient between learning curves of two areas during cue (top), pre (middle) and stim (bottom) periods.

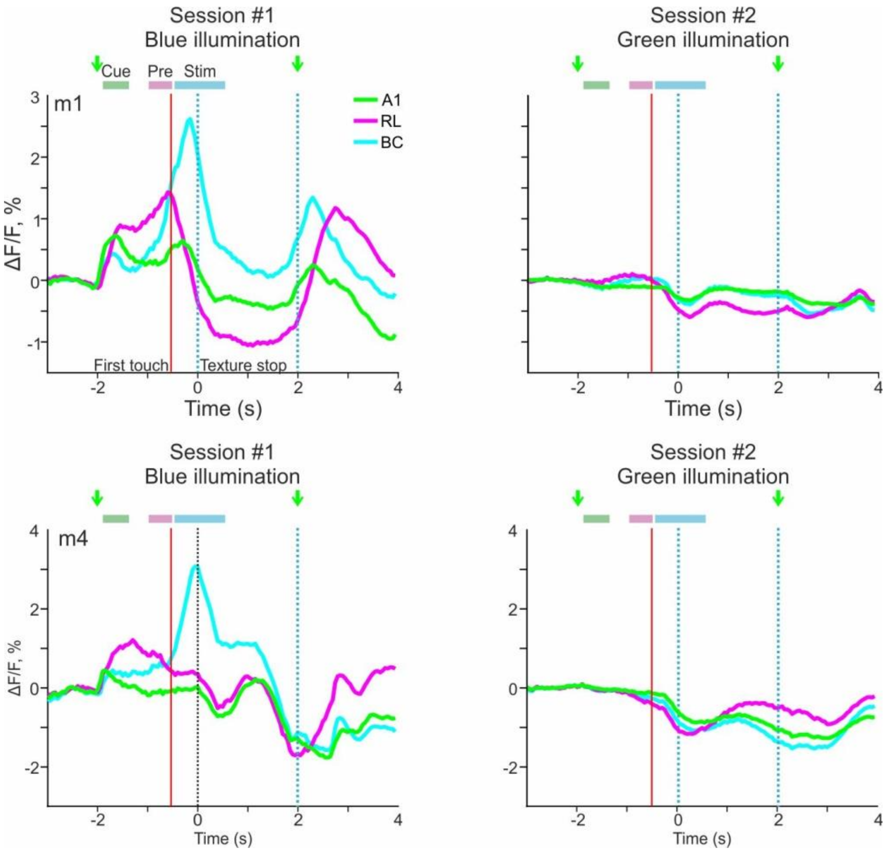


**Supplementary Figure 9 │ Controls for non-calcium related optical signals.** In two mice that reached expert level, we controlled for non-calcium related signal by exciting the wide-field preparation with green light, which is more related to hemodynamic signals. Displayed are the responses in BC, RL and A1 with green light (session #2) compared to the normal excitation with blue light (session #1). There are no major fluorescence changes in the green control, especially during cue and pre periods.
